# Phosphatidylserine exposure by developing astrocytes initiates microglia-mediated developmental cell death

**DOI:** 10.64898/2026.01.20.700660

**Authors:** Caitlin E. Paisley, Kristina Sakers, Leykashree Nagendren, Xi Chen, Francesca Mazzoni, Silvia C. Finnemann, Cagla Eroglu, Jeremy N. Kay

**Affiliations:** Department of Neurobiology, Duke University School of Medicine, Durham, NC 27710, USA; Department of Ophthalmology, Duke University School of Medicine, Durham, NC 27710, USA; Department of Cell Biology, Duke University School of Medicine, Durham, NC 27710, USA; Howard Hughes Medical Institute, Duke University Medical Center, Durham, NC 27710, USA; Department of Biological Sciences, Fordham University, Bronx, NY 10458 USA; Duke Institute for Brain Science (DIBS), Durham, NC 27710, USA

## Abstract

Developmental cell death is classically attributed to apoptosis, yet in mammalian retina, large numbers of developing astrocytes die non-apoptotically during a defined developmental window. Astrocyte death is important for patterning a cellular template that guides angiogenesis, but the underlying mechanism remains unknown. Here we show that healthy developing astrocytes initiate their own elimination by recruiting microglia via regulated exposure of the membrane lipid phosphatidylserine. Experimentally increasing phosphatidylserine exposure in astrocytes, but not neurons, accelerates their removal by microglia without changing how many astrocytes ultimately survive. This acceleration causes profound vascular defects resembling pathological features of retinopathy of prematurity. Genetic disruption of MFGE8, a phosphatidylserine-binding protein, suppresses microglia-mediated astrocyte killing and prevents vascular pathology despite continued phosphatidylserine exposure. This mechanism extends beyond the retina, because phosphatidylserine also initiates astrocyte death in developing cerebral cortex. Together, these findings identify phosphatidylserine exposure as a developmental signal that times microglia-mediated astrocyte elimination, with essential consequences for neurovascular development.

## INTRODUCTION

Astrocytes have essential roles in maintaining the health and function of the central nervous system (CNS), many of which require cell-cell interactions with surrounding structures (Duffy & Eyo, 2025; Raghunathan & Eroglu, 2025; Williamson et al., 2025). Interactions with vasculature, neuron cell bodies, synapses, and even other astrocytes are facilitated by the spatial arrangement and patterning of astrocytes, as established during development. Thus, the developmental mechanisms that control astrocyte patterning are central to establishing mature astrocyte functions (Paisley & Kay, 2021). However, much remains to be learned about astrocyte pattern formation.

One of the main mechanisms for developmental pattern formation is naturally occurring cell death. The impact of death upon astrocyte patterning is particularly well established in the retina, where astrocytes of the retinal nerve fiber layer (RNFL; Figure 1A) are eliminated in large numbers during a specific developmental epoch – the first two postnatal weeks in mice. At the end of the developmental death window, astrocyte numbers have been reduced by three-fold (Paisley & Kay, 2021; Puñal et al., 2019). This massive culling exerts a dramatic effect on the network of astrocyte arbors that fills the RNFL. To date, key mechanisms underlying this large-scale death remain unresolved. First, unlike most instances of naturally occurring cell death which are carried out via apoptosis (Barres et al., 1992; Cecconi et al., 1998; Raven et al., 2003; Suzanne & Steller, 2013; Yamaguchi & Miura, 2015), RNFL astrocytes die via a different death mechanism involving microglia, which we have termed death by phagocyte (Puñal et al., 2019). While microglia are responsible for astrocyte death, the sequence of events that initiates death remains unknown. Without this information, it remains unclear how healthy astrocytes are selected to die. Second, it is not known how the timing of the death period is determined. The factors that open and close the death window are critical determinants of how many astrocytes will ultimately survive to maturity; therefore, identification of such factors will enable insight into the developmental control of astrocyte numbers. Third, it is not yet clear whether this death mechanism is specific to the retina, or if it reflects a general mechanism for developmental astrocyte elimination.

**Figure 1:**
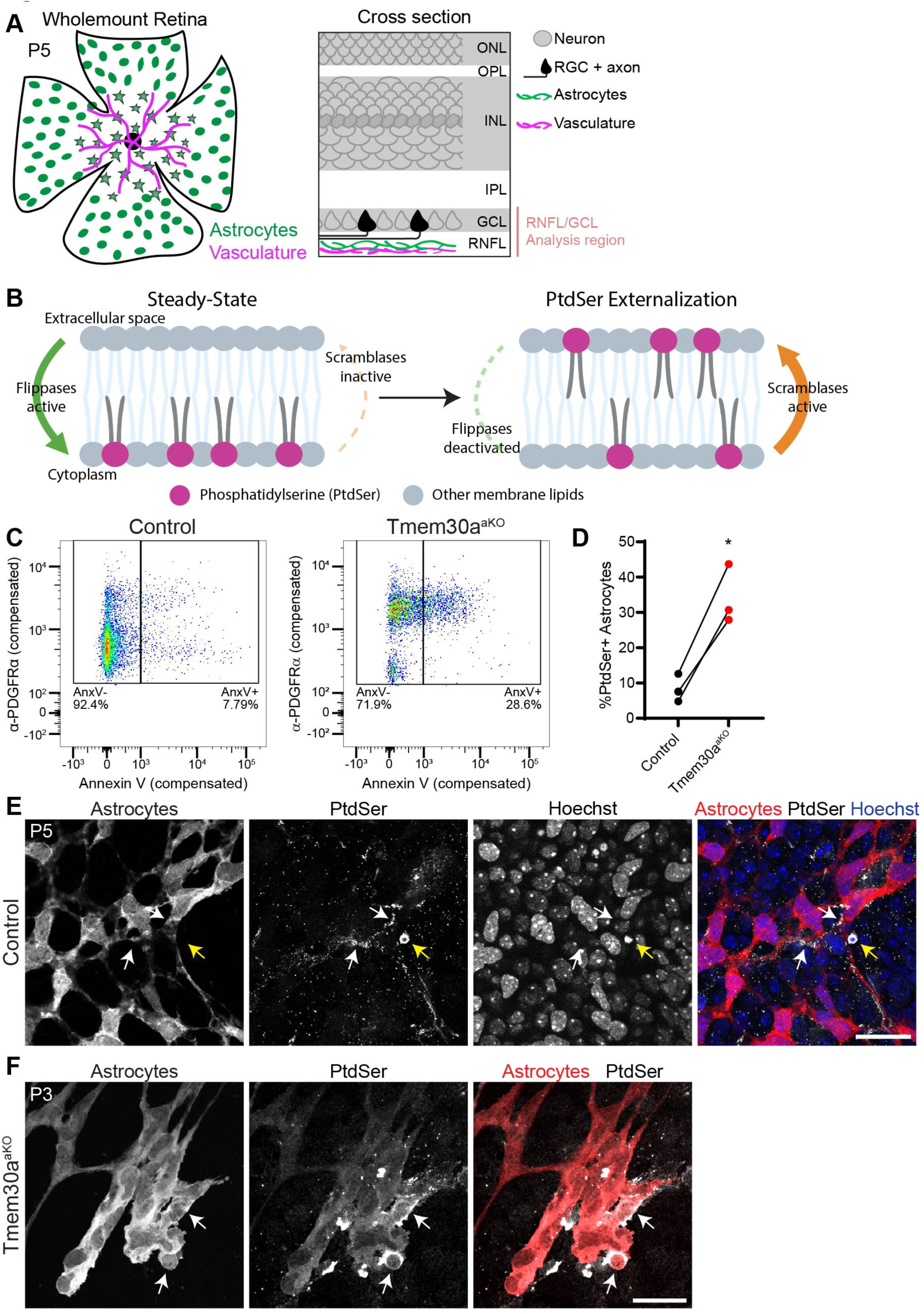
Astrocytes expose cell-surface phosphatidylserine during their developmental death period. A) Illustrations depicting mouse retina and its layers. Left, *en face* view of P5 mouse retina focusing on the retinal nerve fiber layer (RNFL). At this age, astrocytes are found throughout the RNFL, whereas vessels have colonized approximately half of the retina, growing centrifugally from the optic nerve head (center). Astrocytes mature as vessels pass over them (stars, mature astrocytes; ovals, astrocyte precursors in avascular retina). Right, illustration depicting a retinal cross-section. Astrocytes and vasculature (as in B) are found in the RNFL. ONL/INL, outer/inner nuclear layer; OPL/IPL, outer/inner plexiform layer; GCL, ganglion cell layer. Red bar indicates depth of confocal Z-stacks acquired for E, F, and other *en face* confocal images throughout the study. B) Diagram depicting PtdSer localization within the plasma membrane and its control by flippases and scramblases. At steady-state (left), PtdSer is confined to the inner leaflet by flippase activity, which removes PtdSer from the outer leaflet. Signaling events initiated in specific cellular contexts, such as apoptosis or platelet activation, cause scramblase activity to predominate over flippase activity, favoring PtdSer translocation to the outer leaflet where it is exposed on the cell surface. C) Flow cytometry plots for a representative P5 litter showing fraction of PDGFRα+ retinal astrocytes labeled with Annexin V, a marker of PtdSer exposure. Left, control mice; right, *Tmem30a^aKO^*mutants. Plots show only those cells passing the PDGFRα gate; full gating strategy shown in Supplemental Figure 1. D) Graph summarizing flow cytometry results (as in C) for three separate litters containing control and *Tmem30a^aKO^* animals. The fraction of Annexin+ astrocytes increased by ∼25% in *Tmem30a^aKO^* animals compared to controls. Lines indicate samples from the same litter. Statistics: two-tailed, paired *t* test, \**p* = 0.014. E,F) Live staining of wholemount retinas with anti-PtdSer to reveal externalized PtdSer. Astrocytes labeled by genetically encoded tdTomato (red, GFAP-cre; *Rosa26^Ai14^*). E, Control retina; F, *Tmem30a^aKO^*retina. Astrocytes exposing cell-surface PtdSer are marked by white arrows. Yellow arrow, PtdSer+ apoptotic cell with characteristic pyknotic nuclear morphology. Blue, Hoechst nuclear counterstain. Scale bars: 25 µm. B, modified from Paisley & Kay, 2021.

It is important to resolve these questions because, through its impact on RNFL astrocyte patterning, astrocyte death is poised to have a major impact on visual development and function. This impact is due to the fact that RNFL astrocytes are the primary drivers of retinal angiogenesis. Astrocytes exert their pro-angiogenic effects both by recruiting vessels into the tissue and by imposing spatial pattern onto the developing RNFL capillaries (Gerhardt et al., 2003; Osborne et al., 2004; Stone et al., 1995). Each of these astrocytic functions is severely perturbed by developmental alterations to astrocyte numbers: Both overproduction (Duan & Fong, 2019; Fruttiger et al., 1996; Perelli et al., 2021) and underproduction (Derbyshire et al., 2023; O’Sullivan et al., 2017; Tao & Zhang, 2016) of astrocytes can cause major deleterious effects on the pattern of the astrocyte network and the associated vasculature, highlighting the need for developmental mechanisms that precisely control astrocyte numbers. Furthermore, many of the vessel phenotypes resulting from changes in astrocyte density are similar to the pathologies seen in retinopathy of prematurity (ROP), a human disorder of retinal vascular development and a leading cause of preventable childhood blindness worldwide (Lucchesi et al., 2022). Therefore, dysregulation of astrocyte death could be relevant to ROP pathology.

To understand how astrocyte death is initiated, we sought to identify molecular cues that trigger their killing by microglia. We focused on astrocyte-derived signals, because we previously observed a striking correlation between astrocyte numbers and microglial engulfment behavior: As astrocyte numbers increased during the first postnatal week, due to proliferation and migration into the retina, the frequency with which microglia ingested astrocytes increased in lockstep. Subsequently, as astrocyte numbers declined due to ongoing death, microglial engulfment activity also declined (Puñal et al., 2019). This link between astrocyte numbers and microglial phagocytic activity suggests that astrocytes themselves are the source of molecular cues that recruit microglia to kill them.

Here we investigated phosphatidylserine (PtdSer) as a potential signal for the recruitment of microglia and initiation of astrocyte death. PtdSer, a membrane lipid, is normally confined to the inner leaflet of the plasma membrane due to the activity of phospholipid flippases. However, upon activation of lipid scramblases, PtdSer can be externalized to the outer leaflet to initiate cell-cell signaling (Figure 1B). Externalized PtdSer has been particularly well characterized as an “eat-me” signal inducing macrophage-mediated engulfment (Segawa & Nagata, 2015). As such, cell-surface PtdSer is present in many situations that require phagocytic clearance, including apoptosis and synapse elimination (Li et al., 2020; Park et al., 2021; Scott-Hewitt et al., 2020; Segawa & Nagata, 2015). In this eat-me signaling context, PtdSer can be bound by several different extracellular opsonins which can act redundantly with each other; these include milk fat globule-EGF factor 8 (MFGE8), Gas6, Protein S, and C1q proteins. The opsonins, in turn, bind to receptors expressed by phagocytes to promote engulfment (Fricker et al., 2012; Gaboriaud et al., 2011; Lemke, 2013; Nagata, 2018). While best known as an eat-me signal, PtdSer externalization also occurs in a variety of other transcellular signaling contexts (Segawa & Nagata, 2015). Thus, in addition to debris removal, it is plausible that the phagocyte recruitment machinery controlled by PtdSer could instead serve as a trigger for the removal of live, healthy astrocytes during their death window.

To test this hypothesis, we employed a mouse genetic strategy to manipulate PtdSer externalization. At steady state, PtdSer is absent from the outer membrane leaflet because of high constitutive flippase activity (Figure 1B); however, when flippases are experimentally inhibited, PtdSer accumulates on the cell surface (Segawa & Nagata, 2015). Here, we genetically inactivated TMEM30A, a key flippase subunit, to abrogate flippase activity in a cell-type-specific manner, and confirmed prior reports that this manipulation increases PtdSer externalization. We found that forced PtdSer exposure in retinal astrocytes, but not in neurons, drives microglial recruitment and accelerates death without changing the overall number of astrocytes that die. This change to the kinetics of astrocyte elimination caused severe vascular defects, some resembling ROP pathologies. We further identified a molecular pathway, mediated by MFGE8 signaling, required for microglia recruitment and astrocyte killing. Blocking this pathway substantially rescued the astrocyte and vascular defects caused by excess PtdSer exposure. Finally, we showed that this PtdSer-mediated death mechanism operates not only in retinal astrocytes, but also in astrocytes of the cerebral cortex. Altogether, our findings highlight how PtdSer can serve as a “kill-me” signal during astrocyte pattern formation, with major functional consequences for CNS development.

## RESULTS

### Astrocytes expose cell-surface PtdSer during their developmental death window

To investigate the possibility that PtdSer might be a cell-surface signal that promotes astrocyte death, we first assayed for its presence on the surface of developing retinal astrocytes (Figure 1A) at the time when they are dying. The period of astrocyte death in mice extends from shortly after birth until postnatal day (P) 14, with a peak of astrocyte loss occurring at P5 (Puñal et al., 2019). Therefore, we examined PtdSer externalization in P3-P6 mice using two different PtdSer-binding probes. Importantly, because these probes are cell impermeant, they selectively label cell-surface PtdSer when applied to live cells or tissues. First, we used the well-characterized PtdSer probe Annexin V to quantify PtdSer exposure using flow cytometry. By co-labeling with an astrocyte-specific marker (anti-PDGFRα), we determined that 8.3% ± 2.3% of astrocytes expressed cell-surface PtdSer at P5-P6 (Figure 1C,D; Supplemental Figure 1). This finding suggests that developing astrocytes do indeed expose PtdSer, but only a subset of them do so at any given time.

Second, to examine the morphology of PtdSer+ astrocytes in situ, a monoclonal antibody against PtdSer was used to label live retinal explants, as previously described (Ruggiero et al., 2012). Labeled retinas were subsequently fixed and stained for astrocyte markers, to determine whether anti-PtdSer had bound to astrocytes in the live tissue. Consistent with the flow cytometry analysis, this method revealed cell-surface PtdSer labeling on a subset of astrocytes (Figure 1E). PtdSer was also detected on apoptotic neurons with a pyknotic morphology (Figure 1E), confirming the efficacy of the labeling method. Whereas apoptotic (i.e. dying) neurons had a shriveled appearance, PtdSer+ astrocytes were morphologically similar to their PtdSer-negative neighbors, with numerous arbors and normal-sized somata. Thus, PtdSer exposure is not a marker of astrocytes that have already died; rather, it is present on the cell surface of astrocytes that are apparently alive and healthy. Together, these results are consistent with the idea that cell-surface PtdSer exposure could initiate developmental astrocyte death.

### Enhancing astrocyte PtdSer externalization using *Tmem30a^flox^* mutant mice

To test whether PtdSer drives astrocyte death, we used a genetic strategy to manipulate cell-surface PtdSer exposure. *Tmem30a* encodes the obligate beta subunit of the major family of flippases, the P4-ATPases, and is required both for proper flippase trafficking to the plasma membrane and for lipid transport activity (Segawa & Nagata, 2015). Knockout of *Tmem30a* broadly impairs the activity of most P4-ATPases, leading to PtdSer accumulation on the outer membrane leaflet (Segawa et al., 2014). A conditional *Tmem30a^flox^* mutant mouse strain has previously been used in neurons to achieve cell type-specific elevation of PtdSer exposure in vivo (Park et al., 2021; Zhang et al., 2019). We therefore adopted this same strategy to drive PtdSer exposure in astrocytes. An astrocyte-specific *Tmem30a* knockout line (denoted *Tmem30a^aKO^*) was generated by crossing the *Tmem30a^flox^* strain with a GFAP-Cre mouse line that we have characterized extensively (O’Sullivan et al., 2017; Puñal et al., 2019). In this Cre line, the majority of astrocytes undergo Cre recombination by P1 (O’Sullivan et al., 2017), as confirmed by breeding a Cre-dependent fluorescent marker (tdTomato, encoded by the *Rosa26^Ai14^*allele) into the *Tmem30a^aKO^* background (Supplemental Figure 2A,B). Inclusion of the Cre reporter enabled us to investigate the effects of *Tmem30a* knockout on an individual cell level, by discriminating between Cre+ and Cre-astrocytes.

To validate this genetic strategy, we first verified astrocyte expression of the P4-ATPases using single-cell RNA-seq data (Li et al., 2024). At least seven of the fourteen genes encoding P4-ATPases were broadly expressed across mouse retinal cell types, including astrocytes (Supplemental Figure 1C). Therefore, astrocytes likely possess P4-ATPase activity that could be impaired by the loss of *Tmem30a*. Moreover, because astrocytes express so many different P4-ATPases, the *Tmem30a* single knockout is a more efficient strategy for inhibiting flippases than direct knockout of P4-ATPases themselves. Next, to test whether deleting *Tmem30a* indeed increases PtdSer externalization, we subjected mutant astrocytes to the Annexin V flow cytometry assay. This analysis confirmed that the fraction of PtdSer+ astrocytes was substantially higher in P5-P6 *Tmem30a^aKO^* mice than in littermate controls (Figure 1C,D). Live staining at P3 and P5 for anti-PtdSer also showed stronger labeling for externalized PtdSer on astrocytes (Figure 1F). Taken together, these analyses show that *Tmem30a^aKO^* has the desired effect of increasing PtdSer exposure by retinal astrocytes.

### Forced PtdSer exposure alters astrocyte morphology and accelerates their death

To assess how astrocyte development is impacted by enhanced PtdSer exposure, we first examined astrocyte morphology and numbers at P5 and P10 in *Tmem30a^aKO^* mice. Two different astrocyte markers were used: Sox9, which labels astrocyte nuclei; and *Ai14-*tdTomato driven by the GFAP-Cre transgene (denoted GFAP-tdT), which reveals astrocyte somata and processes. Comparing *Tmem30a^aKO^* mutants and their littermate controls at P5, we noted striking anatomical changes to the astrocyte population: Astrocyte density was greatly reduced relative to littermate controls; and the remaining mutant tdT+ astrocytes were smaller, had fewer processes, and showed fewer contacts with their neighbors (Figure 2A,B). The mutant phenotype was similar at P10, with an even more obvious loss of arbors from remaining astrocytes (Figure 2C,D). Additionally, we also noted that mutant astrocytes tended to aggregate, in a manner that was never observed in controls. These aggregates of tdT+ mutant cells formed at specific retinal locations, which varied with age: At P5, aggregates were selectively localized to the vascular wavefront in the middle of the retina (see Figure 1A), while at P10, tightly packed astrocyte clumps were observed in the retinal periphery (Figure 2E). Together, these observations indicate that excess PtdSer exposure is associated with multiple anatomical features that would be expected if astrocytes are dying – i.e., loss of morphological complexity, cell shrinkage, and decreased cell density.

**Figure 2:**
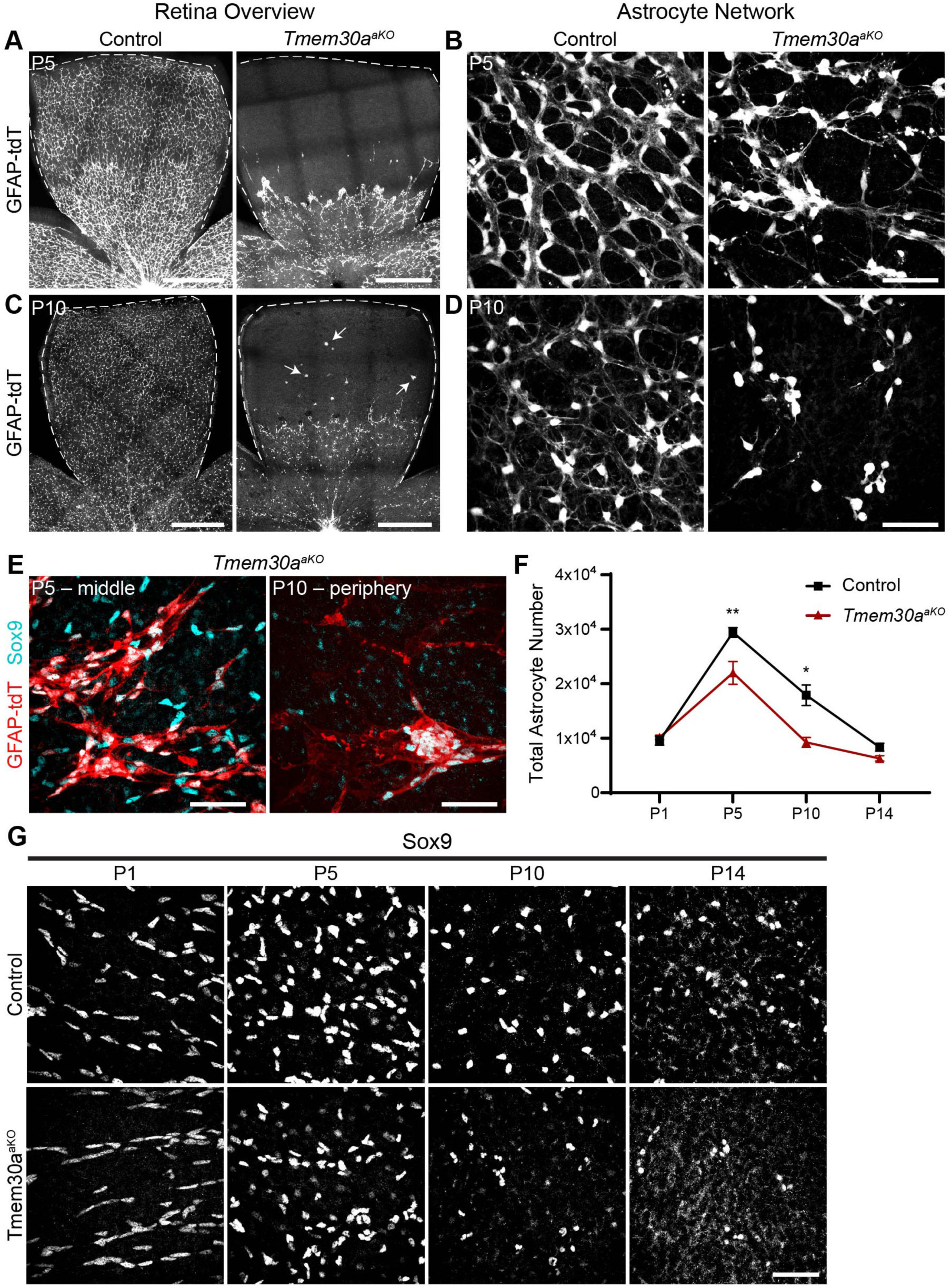
Forced cell-surface PtdSer exposure accelerates astrocyte death. A-D) Retinal astrocyte morphology at P5 (A,B), and P10 (C,D) in control and *Tmem30a^aKO^* animals. A,C: Stitched confocal tiles showing single lobes of wholemounted retinas. Note substantial loss of astrocytes from *Tmem30a^aKO^*retinas. Arrows, astrocyte soma aggregates in peripheral mutant retina. B,D: high-power images showing astrocyte arbors. *Tmem30a^aKO^* astrocyte arbors are less ramified and not connected in a network as they are in control animals. Astrocytes labeled by genetically encoded tdTomato driven by GFAP-cre (GFAP-tdT). E) Examples of astrocyte aggregation in *Tmem30a^aKO^* retinas. At P5 (left), astrocytes form aggregates near the vessel wavefront in middle retina. At P10 (right), larger clumps with more densely packed astrocytes form in the periphery of the retina. All of the astrocytes in these aggregates are Cre+ (GFAP-tdT+, red), compared to astrocytes outside of the aggregates that are mostly Cre– (only Sox9+, cyan). F) Total astrocyte numbers across development in control and *Tmem30a^aKO^* animals. Total number estimates based on eccentricity-weighted density measurements and retina area (see Methods). Significant decreases in astrocyte number in mutants at P5 and P10. Statistics: Multiple *t* tests with Bonferroni-Dunn correction for multiple comparisons. P5 \*\**p* = 0.007, P10 \**p* = 0.025. Sample sizes ≥ 3 animals per timepoint and genotype. Error bars: mean ± SEM. G) Representative images of Sox9-labeled astrocyte nuclei at different developmental timepoints in control and *Tmem30a^aKO^* retinas. P1, middle retina; P5, central retina; P10, middle retina; P14, middle retina. Scale bars: A,C: 500 µm; B,D,E,G: 50 µm.

To evaluate whether these anatomical changes do indeed reflect a loss of cells, we counted Sox9+ astrocytes in retinal wholemounts and estimated total astrocyte numbers as previously described (O’Sullivan et al., 2017; Puñal et al., 2019). For these cell counts and subsequent analyses, the control group included both *Tmem30a* wild-type animals (i.e. two functional gene copies) as well as GFAP-Cre+;*Tmem30a^flox/+^* heterozygotes, because astrocyte phenotypes did not differ between these genotypes (Supplemental Figure 2E). Comparing *Tmem30a^aKO^* mutants to controls, astrocyte numbers were significantly lower in mutants at both P5 and P10, suggesting that elevating PtdSer exposure enhances astrocyte death (Figure 2F,G). This phenotype was not accompanied by an increase in cleaved caspase-3 expression; indeed, with only one exception, every astrocyte we surveyed was cleaved caspase-3-negative (Supplemental Figure 2F,G). This finding suggests that death occurs by the same non-apoptotic mechanism that occurs in wildtype astrocytes (Puñal et al., 2019).

To address how the dynamics of the astrocyte death period are altered by forced PtdSer exposure, we expanded our analysis of astrocyte numbers to additional timepoints (P1 and P14), generating a timecourse that spans the first two postnatal weeks (Figure 2F,G). At P1, we noted that astrocyte migration was somewhat delayed in *Tmem30a^aKO^* mutants; however, there was no difference in astrocyte numbers between mutants and controls (Figure 2F,G; Supplemental Figure 2C,D). Subsequently, during the P1-P5 period when wildtype astrocyte numbers are increasing due to proliferation and migration into the retina (Paisley & Kay, 2021), mutant numbers also increased. However, this increase was blunted in mutants relative to controls, suggesting that the balance between cell addition and removal was shifted by the loss of *Tmem30a*. Astrocyte numbers remained lower in mutants at P10. Surprisingly, however, astrocyte numbers declined only minimally between P10 and P14, such that the number of astrocytes was not significantly different between mutants and controls at the conclusion of the death period (Figure 2F,G). These observations demonstrate that forced PtdSer exposure does not cause excess astrocyte death but instead alters the dynamics of the death period such that death is accelerated.

The lack of excess death strongly suggests that deletion of *Tmem30a* does not kill astrocytes through a nonspecific toxic mechanism. To further rule out the possibility of nonspecific toxic effects, we generated neuron-specific *Tmem30a* knockout animals. Because developing neurons die by apoptosis, which cannot be induced by forced PtdSer exposure (Segawa et al., 2014), we expected minimal effects on neuron survival. Indeed, despite effects on synapse pruning, previous studies in which *Tmem30a* was genetically removed from brain neurons did not document aberrant neuronal loss (Li et al., 2021; Park et al., 2021). To test whether this holds true for retinal neurons, we used *Chat^Cre^* driver mice to delete *Tmem30a* from starburst amacrine cells (SACs), a cholinergic retinal neuron population. In these *Tmem30a^sacKO^* animals, SAC numbers were unchanged compared to littermate controls (Figure 3A,B). Furthermore, in striking contrast to astrocytes which largely lost their fine processes upon PtdSer upregulation, dendritic arborizations of starburst neurons were maintained in *Tmem30a^sacKO^* mice (Figure 3A). These findings indicate that astrocytes and neurons respond differently to increased PtdSer exposure, and that acceleration of developmental cell death by PtdSer is an astrocyte-specific phenomenon.

**Figure 3:**
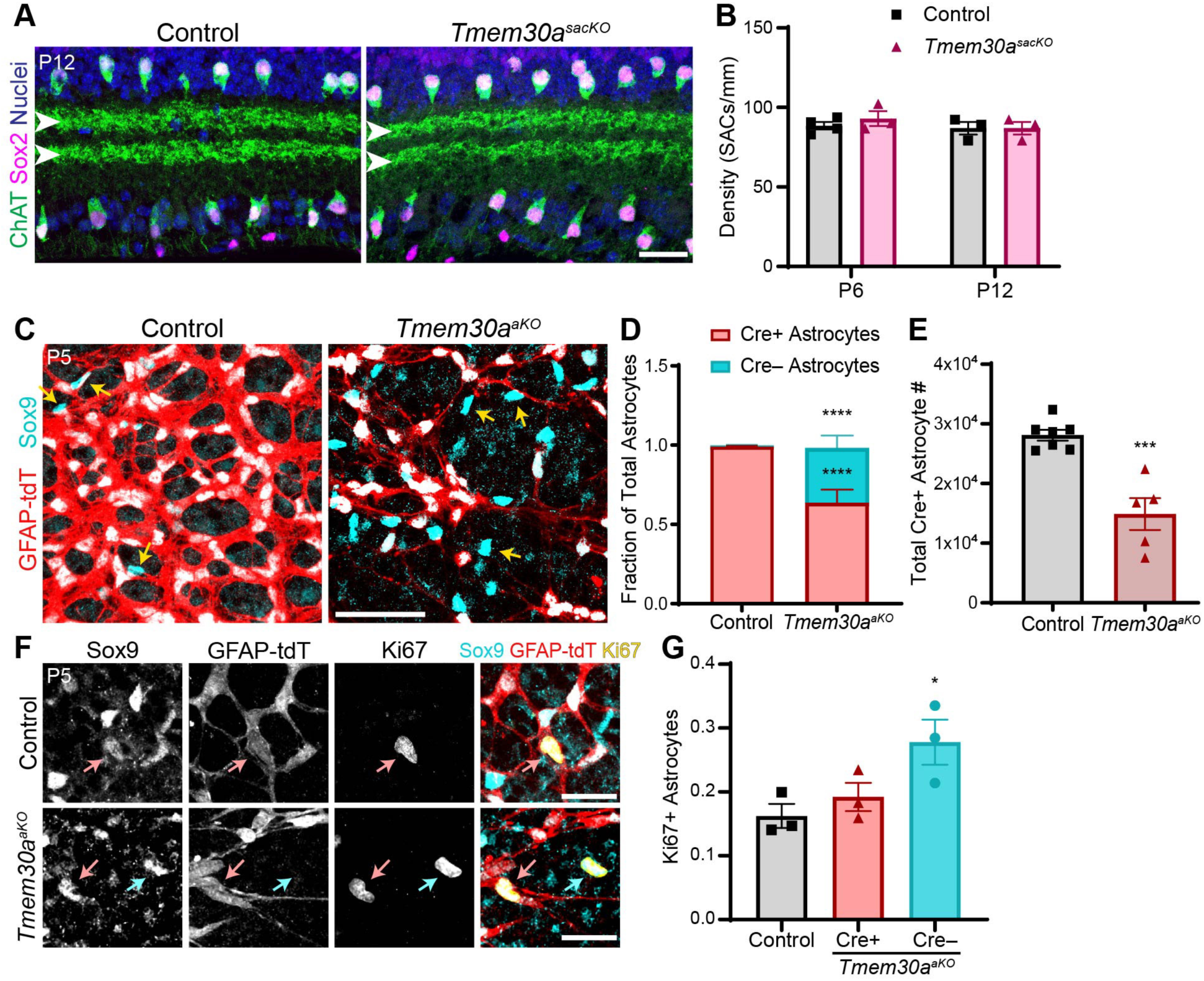
*Tmem30a* death phenotype is astrocyte-specific and cell-autonomous. A,B) Deletion of *Tmem30a* from starburst amacrine cells (*Tmem30a^sacKO^*) does not affect SAC survival or dendritic arborization. A, Representative images of retinal cross-sections from P12 control and *Tmem30a^sacKO^*animals. SACs labeled with ChAT (green, soma and dendrites) and Sox2 (magenta, nuclear marker). SAC dendrites (arrowheads) arborize in sublayers S2 and S4 of the IPL. Blue, Hoechst counterstain. B. Quantification of SAC density at P6 and P12. Statistics: Multiple *t* tests with Bonferroni-Dunn correction for multiple comparisons. P6 *p* = 0.792, P12 *p* > 0.999. Sample sizes shown on graph. C,D) Selective loss of Cre-reporter+ astrocytes in *Tmem30a^aKO^* retinas. C, Representative en face images of P5 retinal wholemounts stained for Sox9 (cyan) and tdTomato Cre reporter driven by GFAP-Cre (GFAP-tdT, red). D) Quantification of Cre+ (Sox9+GFAP-tdT+) and Cre–(Sox9+GFAP-tdT–) astrocytes. In controls, nearly all Sox9+ astrocytes also express GFAP-tdT (98.6%); yellow arrows highlight rare GFAP-tdT– astrocytes. The number of Sox9+ astrocytes without Cre reporter increased markedly in *Tmem30a^aKO^*retina. Statistics: 2-way ANOVA showing a significant interaction between genotype and colocalization (\*\*\*\**p* < 1×10^-7^), with post-hoc Sidak’s test corrected for multiple comparisons. Significant differences between control and *Tmem30a^aKO^* Cre+ astrocyte fraction (\*\*\*\**p* = 7.00 x 10^−6^) and Cre– astrocyte fraction (\*\*\*\**p* = 1.39 x 10^−5^). Sample size: N ≥ 5 for each group. E) Total number of GFAP-tdT+ (i.e. Cre+) astrocytes per retina at P5, estimated based on eccentricity-weighted density measurements (see Methods). Effects on Cre+ astrocyte survival were larger than effects on the full Sox9+ astrocyte population (Figure 2F), suggesting selective elimination of Cre+ astrocytes in *Tmem30a^aKO^* retinas. Statistics: two-tailed *t* test, \*\*\**p* = 1.51 x 10^−4^. Sample sizes shown on graph. F,G) Ki67 expression by control and *Tmem30a^aKO^* mutant astrocytes at P5. F: Images showing examples of Ki67+Sox9+ astrocytes that express GFAP-tdT reporter (Cre+, red arrows) and Ki67+Sox9+ cells that lack Cre reporter (Cre–, cyan arrow). G: Quantification of images similar to F, to determine the fraction of dividing astrocytes (Ki67+Sox9+) in controls and in *Tmem30a^aKO^*astrocytes with or without Cre. (Because so few astrocytes were Cre– in control mice, this population was not considered separately.) Mutant astrocytes that escaped Cre were more proliferative than their Cre+ counterparts in *Tmem30a^aKO^*retinas. Statistics: one-way ANOVA showing a significant group effect (\**p* = 0.049), with post-hoc Tukey’s tests corrected for multiple comparisons. The difference in proliferation levels between control astrocytes and Cre–mutant astrocytes was significant (\**p* = 0.048). No significant difference in proliferation levels between control and Cre+ mutant astrocytes was found (*p* = 0.717). Error bars: mean ± SEM. Scale bars: A,F: 25 µm; C: 50 µm.

### Cell-autonomous astrocyte death induced by PtdSer exposure

If PtdSer exposure is a mechanism that selects which astrocytes will die, we would expect death in *Tmem30a^aKO^* mice should be cell-autonomous: astrocytes that expose excess PtdSer should be more vulnerable to death than those that do not. To test this model, we took advantage of the fact that GFAP-Cre is expressed by most, but not all, astrocytes. At P5 (but not P10; see Supplemental Figure S3), approximately 1.5% of Sox9+ cells in control mice were negative for the GFAP-tdT reporter, indicating that they had escaped Cre activity (Figure 3C,D). If PtdSer exposure promotes death in a cell-autonomous manner, this Cre-negative population should be less susceptible to death in *Tmem30a^aKO^* mice relative to their Cre+ mutant neighbors. Indeed, we found that the Cre-negative population expanded massively to ∼35% of Sox9+ cells in *Tmem30a^aKO^*mice (Figure 3C,D). Not all of this increase can be directly attributed to survival effects: Cre-negative astrocytes retaining *Tmem30a* function also became more proliferative compared to their Cre+ neighbors, perhaps as a homeostatic response to replace dying Cre+ cells. Nevertheless, the excess Cre-negative astrocytes generated in this manner did survive long enough to contribute to the overrepresentation of Cre-negative astrocytes in mutant retina. Overall, these findings strongly suggest that escaping the loss of *Tmem30a* does confer a survival advantage. (Figure 3F,G).

Given these findings, it was apparent that the Sox9 cell counts reported in Figure 2F included a substantial number of unrecombined astrocytes, at least at P5. To obtain a more accurate estimate of how strongly PtdSer exposure drives astrocyte death at this age, we specifically quantified astrocytes that expressed the Cre reporter, excluding tdT-negative cells. With this analysis, the decline in astrocyte numbers between P5 mutant and control retinas was approximately doubled in magnitude: whereas Sox9 counts were 25% lower in mutants, GFAP-tdT+ astrocyte counts were 48% lower (Figure 3E). Together, these findings support the conclusion that cell-surface PtdSer exposure promotes astrocyte death in a cell-autonomous manner.

### Brain astrocytes are subject to PtdSer-mediated death

We next addressed whether the PtdSer-driven death phenomenon generalizes to astrocyte populations outside the retina. Astrocytes of the mouse cerebral cortex decline in density during the first two postnatal weeks, suggesting that they also undergo developmental death (Puñal et al., 2019). To test whether the death of these astrocytes is also driven by PtdSer exposure, we sparsely deleted *Tmem30a* from cortical astrocytes using Postnatal Astrocyte Labeling by Electroporation (PALE) (Stogsdill et al., 2017). A plasmid encoding Cre recombinase was electroporated unilaterally into the dorsomedial-caudal cortex of P0-1 *Tmem30a^flox/flox^* mutants, or *Tmem30a^flox/+^* heterozygotes as littermate controls (Figure 4A,B). This method produces a sparse transfection of radial glial progenitor cells (Figure 4A), which at P0-1 are fate-committed to generating astrocytes (Stogsdill et al., 2017). Inclusion of the *Ai14* tdT reporter allele enabled visualization of Cre+ electroporated cells (denoted PALE-tdT labeling). At P7, transfected radial glial progenitors and their astrocyte progeny were visible throughout the targeted region, which included primary somatosensory and visual cortices as well as surrounding regions (Figure 4A-C). By P14, PALE-tdT+ progenitors were no longer present but cortical astrocytes remained labeled (Figure 4E).

**Figure 4:**
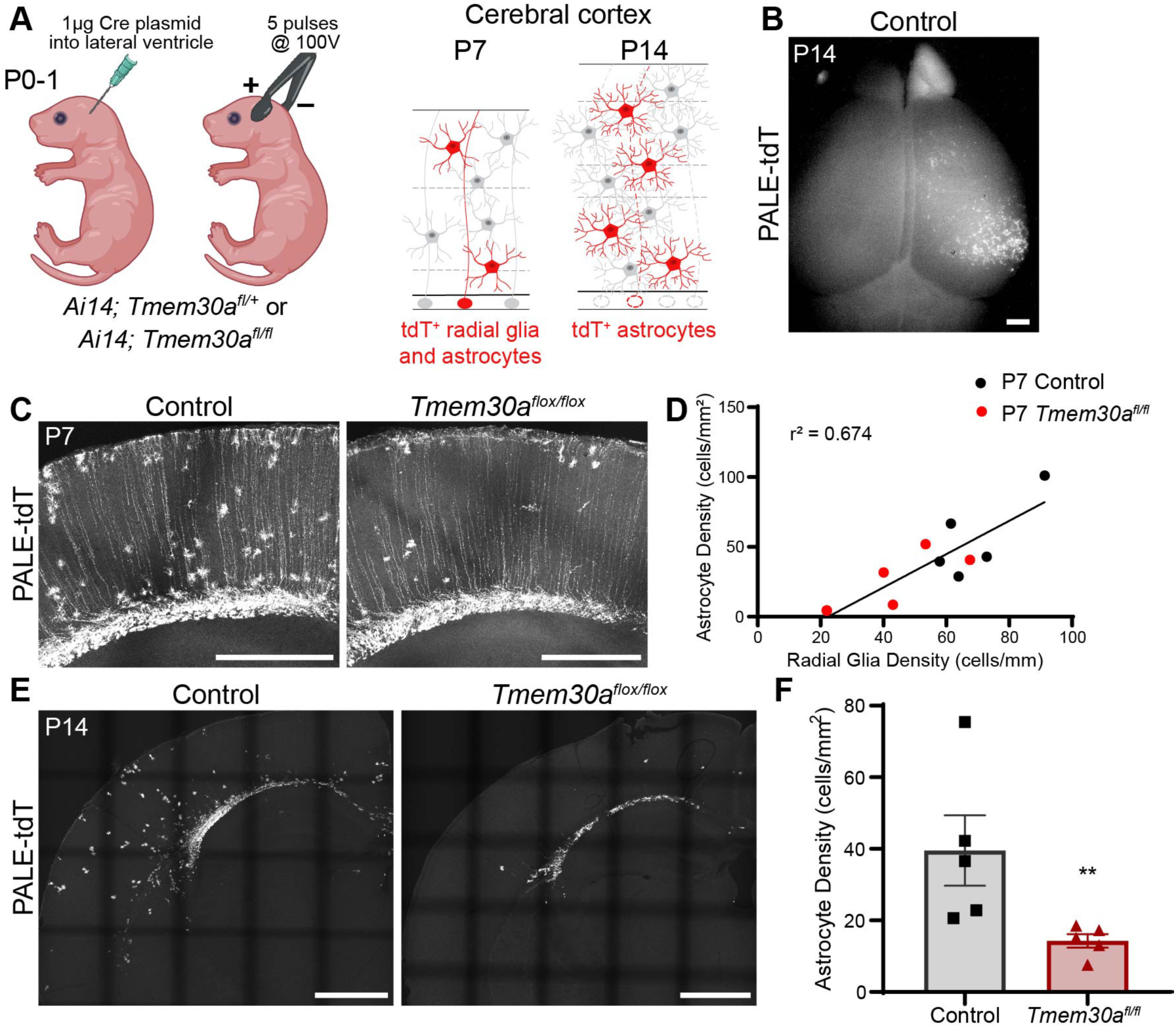
Brain astrocytes are subject to PtdSer-mediated death. A) Overview of postnatal astrocyte labeling by electroporation (PALE). Left, electroporation method schematic, noting plasmids and mouse genotypes used here. *Ai14*, Cre-dependent tdT reporter allele. Right, schematics of cortical cross-sections from *Ai14* PALE mice at specified ages, showing expected pattern of sparse tdT reporter expression (red) by radial glia progenitors and their astrocyte progeny (untransfected cells shown in gray). At P7, both radial glia and astrocytes are tdT+, but by P14 only astrocytes remain visible. B) Widefield epifluorescence photomicrograph showing native tdT fluorescence in the cortex of a control P14 *Ai14* mouse that has undergone PALE with Cre plasmid. C) Representative cross-section images showing PALE-tdT+ radial glia processes in P7 control and *Tmem30a^flox/flox^* cortex. Both radial glia (cells spanning the tissue) and astrocytes (round bushy cells) are visible. D) Scatterplot showing the correlation between density of PALE-labeled radial glia and astrocytes within individual P7 mice. Black dots, control animals; red dots, *Tmem30a^flox/flox^* animals. The efficiency of radial glia transfection varied across individual animals, but there was a consistent relationship between the number of transfected radial glia and the number of labeled astrocyte progeny at P7. This relationship was similar between genotypes, suggesting that loss of *Tmem30a* did not disrupt astrocyte fate specification or differentiation. E) Representative images of PALE-tdT labeled astrocytes in P14 control and *Tmem30a^flox/flox^* cortex, after transfection with Cre plasmid. F) Quantification of PALE-tdT+ astrocyte density from P14 sections similar to E. Astrocyte density was significantly reduced in *Tmem30a^flox/flox^* animals. Statistics: two-tailed Mann-Whitney test, \*\**p* = 0.008. Error bars: mean ± SEM. Scale bars: B,E: 1 mm; C: 500 µm.

To determine the influence of PtdSer exposure on astrocyte survival, we compared Cre-electroporated *Tmem30a^flox/flox^* mutants to littermate controls. If astrocyte survival is affected, PALE-tdT+ astrocyte numbers should be lower in mutants; but because astrocytes derive from radial glia, such a phenotype could also result from decreased radial glia labeling. Therefore, we first counted radial glia in mutants and controls at P7. We did note a trend toward fewer PALE-tdT+ radial glia in *Tmem30a^flox/flox^*animals, suggesting that either electroporation efficiency was coincidentally lower in this group or that radial glial survival was affected (Figure 4D, Supplemental Figure 4A,B). Regardless of the reason for this trend, we observed a consistent relationship between the number of transfected radial glia and the number of astrocyte progeny, which remained similar between control and *Tmem30a^flox/flox^* groups, suggesting that the ability of radial glia to generate astrocytes was not impaired in mutants (Figure 4D). To assess survival of these astrocytes, PALE-tdT+ cells were counted at P14. Strikingly, the density of PALE-labeled astrocytes in *Tmem30a^flox/flox^* mutants was reduced by greater than two-fold (Figure 4E,F). These observations indicate that PtdSer externalization is sufficient to cause developmental death of cortical astrocytes.

To further probe the cell type specificity of this death phenomenon, we used *Olig2^Cre^* to delete *Tmem30a* from the oligodendrocyte lineage (*Tmem30a^opcKO^* mice; Supplemental Figure 4C). Oligodendrocyte precursor cells (OPCs) of the mouse corpus callosum have been shown to undergo substantial developmental death during the first postnatal week (Nemes-Baran et al., 2020). However, our analysis of *Tmem30a^opcKO^* mice suggests that OPC death is not influenced by forced PtdSer exposure, as no effect on callosal OPC morphology or cell density was observed at P7 (Supplemental Figure 4C,D). Altogether, these studies support the idea that developing astrocytes in multiple CNS regions are subject to PtdSer-mediated death, whereas developmental death of other CNS cell types occurs by distinct mechanisms that cannot be induced by PtdSer exposure.

### Accelerated astrocyte death disrupts angiogenesis

To ascertain whether PtdSer-driven death is important for astrocyte development, we next investigated the consequences for astrocyte function when death is dysregulated. For these studies we focused on retinal astrocytes, which have a central role in promoting and patterning retinal angiogenesis (Paisley & Kay, 2021). Developing astrocytes form a physical template that guides growing endothelial cells; patterning of this template can be perturbed by experimental manipulations that either increase or decrease astrocyte numbers (Dorrell et al., 2002; Gerhardt et al., 2003; O’Sullivan et al., 2017; Perelli et al., 2021; Tao & Zhang, 2016). While overall astrocyte numbers at the end of the death period are not altered by astrocyte-specific knockout of *Tmem30a*, this mutation does alter the dynamics of the death period in a manner that could plausibly disrupt template function (Figure 2F). Thus, we investigated whether formation of the astrocyte template and development of the retinal vasculature are disrupted in *Tmem30a^aKO^*mice.

To evaluate the anatomy of the astrocyte arbor network that constitutes the angiogenic template, we stained P10 wholemount retinas for the astrocyte-specific marker GFAP. This analysis revealed that the astrocyte network was severely disrupted in *Tmem30a^aKO^* retinas. Consistent with our observations using the GFAP-tdT reporter (Figure 2A-D), astrocytes were less ramified and less connected to each other (Figure 5A). As a result of this phenotype, as well as the lower absolute astrocyte numbers at this age (Figure 2F,G), retinal coverage by GFAP arbors was reduced nearly 4-fold in *Tmem30a^aKO^* retinas as compared to controls (Figure 5B).

**Figure 5:**
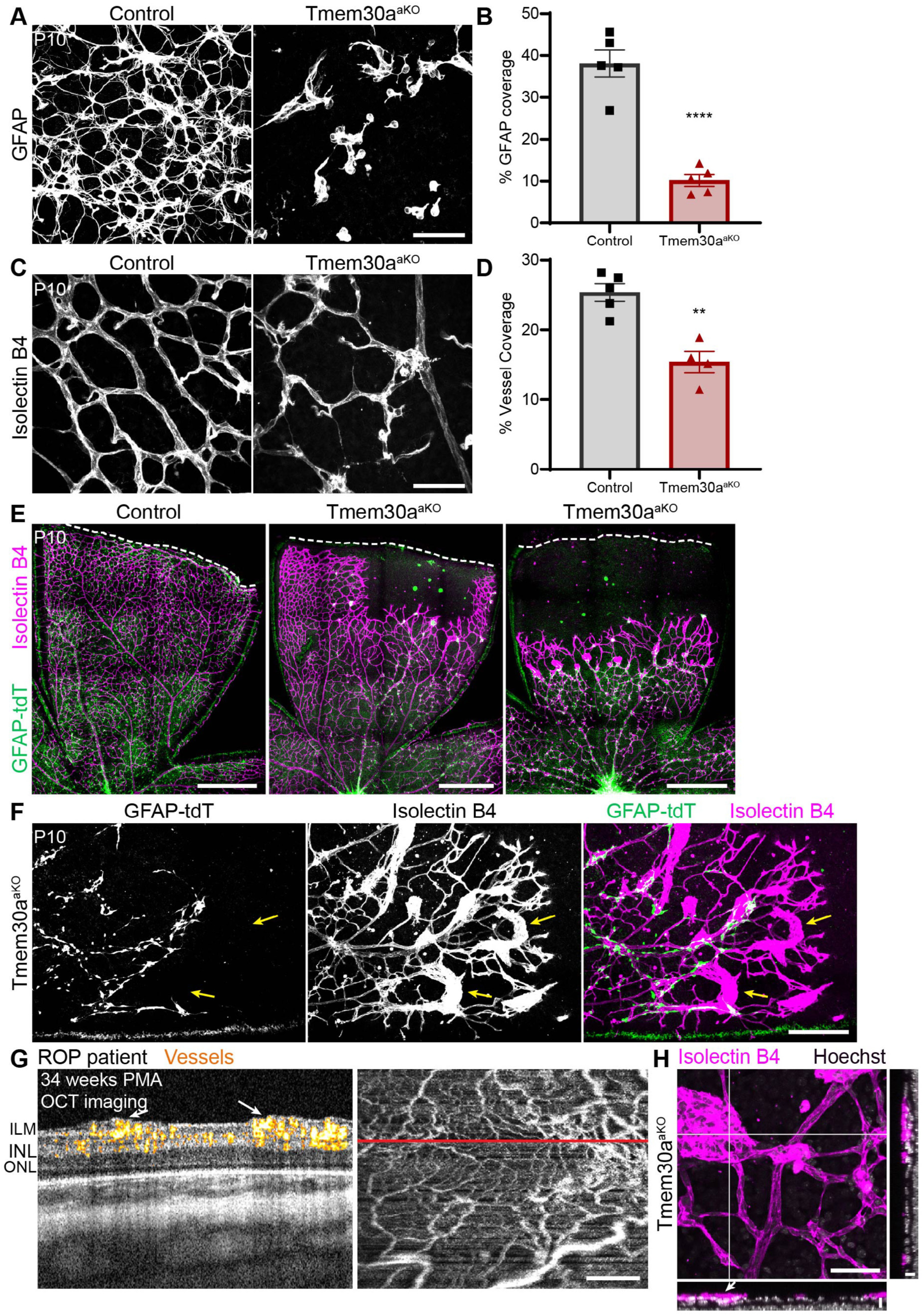
Accelerated astrocyte death disrupts angiogenesis. A,B) The mesh-like network of retinal astrocytes and their arbors is severely disrupted in *Tmem30a^aKO^* mice. A: Representative images showing GFAP coverage of a square field of view (212 µm/side) in the middle region of P10 control and *Tmem30a^aKO^* retinas. B: Quantification of GFAP coverage. Graph shows average coverage from a series of images similar to A, collected to sample uniformly across the entire retina. Statistics: two-tailed *t* test, \*\*\*\**p* = 4.53 x 10^−5^. C,D) Diminished vessel coverage in *Tmem30a^aKO^* retinas. C: Representative vessel images (Isolectin B4 labeling) from the middle region of P10 control and *Tmem30a^aKO^* retinas. D: Quantification of vessel coverage. Graph shows average coverage from a series of images similar to C, collected to sample uniformly across the entire retina. Statistics: two-tailed *t* test, \*\**p* = 0.002. E) Vessels do not cover the entire *Tmem30a^aKO^* retina at P10. Left, control retina stained for vessels (Isolectin B4, magenta) and astrocytes (GFAP-tdT, green), imaged at RNFL level to show primary vascular plexus. Two example *Tmem30a^aKO^* retinas are shown (center, right) illustrating the range of phenotypes. The milder *Tmem30a^aKO^* example (center) illustrates how vessels fail to enter regions where the only remaining astrocytes are confined to aggregates. The severe example (right) is more representative of the typical mutant retina. Even in regions that are vascularized, the mutant vascular network is less dense, as seen in the central and middle regions of both *Tmem30a^aKO^* lobes. Dashed lines show the edge of each retina. F) Higher magnification view of the P10 *Tmem30a^aKO^* vessel wavefront, stained as in E, showing hypertrophic vascular tufts (yellow arrows). Vessels (magenta) typically form these structures when they lack underlying astrocytes (green). G) Bedside handheld optical coherence tomography angiography (OCTA) images of retinal vasculature in eye with neovascularization from previously published data set (Chen et al., 2024). Images taken at 34 weeks post menstrual age from a preterm infant who was born at 23 weeks gestational age; imaging session was 1 day prior to treatment for retinopathy of prematurity (ROP). Left: cross-sectional B-scan with flow overlay at the site of the red line. Scan shows prominent vascular flow and mild elevation of the inner retinal surface (arrows). Right: *en face* image of vascular pattern. OCTA flow signal appears yellow and red in cross-section. Signal is summed across the volume to create the white vascular pattern of the *en face* image. H) Example image of a large hypertrophic vascular tuft in P10 *Tmem30a^aKO^* retina. Vessels (Isolectin B4, magenta) extend further into the underlying layers of retinal cells (arrow) similar to the expansion of vessel width seen in infant ROP imaging (G). White lines show site of orthogonal views. Hoechst counterstain (gray) to show perturbation of underlying retinal cell layers in cross-sections. *En face* view of vasculature shows a single optical z-plane. Error bars: mean ± SEM. Scale bars: A,C,H: 50 µm; E,G: 500 µm; F: 200 µm; H (orthogonal views): 10 µm.

Given these major changes to the astrocyte template, we surmised that angiogenesis might also be impaired. Indeed, quantification of retinal vascular coverage revealed a major reduction in *Tmem30a^aKO^* mutants (Figure 5C,D). One reason for this diminished vessel coverage was a failure of vessels to enter peripheral retinal regions, where the astrocyte phenotype was most severe and where, in many cases, the template was entirely absent (Figure 5E). These findings indicate that dysregulation of astrocyte death period timing results in the elimination of the angiogenic template from peripheral retina before vessels arrive, thereby preventing angiogenesis.

Additionally, even within central regions that contained both astrocytes and vessels, we observed vascular anomalies. The most prominent anomaly was the frequent presence of hypertrophic vascular tufts at the vascular wavefront, which were reminiscent of the pathological intraretinal neovascularization seen at the vascular wavefront in preterm infants with severe ROP (Figure 5E,F) (Lucchesi et al., 2022). A close examination of the tufts in mutant retina revealed that, in striking contrast to the normal strict pairing of astrocytes and vasculature, the hypertrophic tufts lacked underlying astrocytes – perhaps due to local death after vessels had arrived, or perhaps due to attempted sprouting into regions locally lacking astrocytes (Figure 5F). This observation suggests that loss of astrocytic support might contribute to certain aspects of ROP disease pathology. To better compare the mouse and human vascular phenotypes, we examined an existing dataset of handheld optical coherence tomography angiography (OCTA) images, collected with an investigational handheld OCTA device for a study on neovascular phenotypes in infants with severe ROP (Chen et al., 2024). These OCTA images revealed that ROP patients have two types of neovascularizations. The first type is extraretinal, involving vessels that extend out of the retina into the vitreous; such tufts were not observed in *Tmem30a^aKO^* mice. However, ROP patients also exhibit intraretinal neovascularization, which grows within the RNFL and displaces neuronal layers beneath it (Figure 5G, left). We noted striking anatomical similarities between these intraretinal neovascularizations and those observed in *Tmem30a^aKO^* mice – notably the displacement of underlying neuronal layers (Figure 5G,H). Together, these studies demonstrate that vessel development is severely perturbed by dysregulation of the PtdSer-driven astrocyte death mechanism, highlighting the importance of cell-surface PtdSer signaling not only for astrocyte development but also the retina as a whole.

### Microglia are recruited to PtdSer-expressing astrocytes

We next sought to determine the molecular and cellular mechanism by which PtdSer exposure leads to astrocyte death. Our previous work showed that microglia are directly involved in the developmental death of retinal astrocytes such that, when microglia are experimentally removed, fewer astrocytes die (Puñal et al., 2019). Thus, we hypothesized that microglia would also mediate astrocyte death induced by forced PtdSer exposure. If this is true, we would expect microglia to be recruited to engulf astrocytes in *Tmem30a^aKO^* mice. Supporting this idea, we observed swarms of microglia in mutant retina that were selectively associated with GFAP-tdT+ astrocyte aggregates (Figure 6A). This swarming behavior altered microglia distributions in two different ways. First, it led to a significant increase in total microglia number within the RNFL and GCL of *Tmem30a^aKO^* retinas compared to controls (Figure 6B,C). Second, the planar distribution of microglia within the RNFL was altered in *Tmem30a^aKO^*mutants: Instead of being evenly spaced across the RNFL, as was observed in control retinas, microglia in *Tmem30a^aKO^* retinas accumulated at astrocyte aggregates while leaving adjacent RNFL regions nearly barren (Figure 6A,B). Microglia that were recruited to astrocyte-rich regions of mutant retina frequently contained GFAP-tdT+ astrocyte debris, indicating that recruitment was accompanied by astrocyte engulfment (Figure 6D). Furthermore, live staining for PtdSer revealed that the sites where microglia contacted mutant astrocytes often corresponded to sites of PtdSer exposure (Figure 6E).

**Figure 6:**
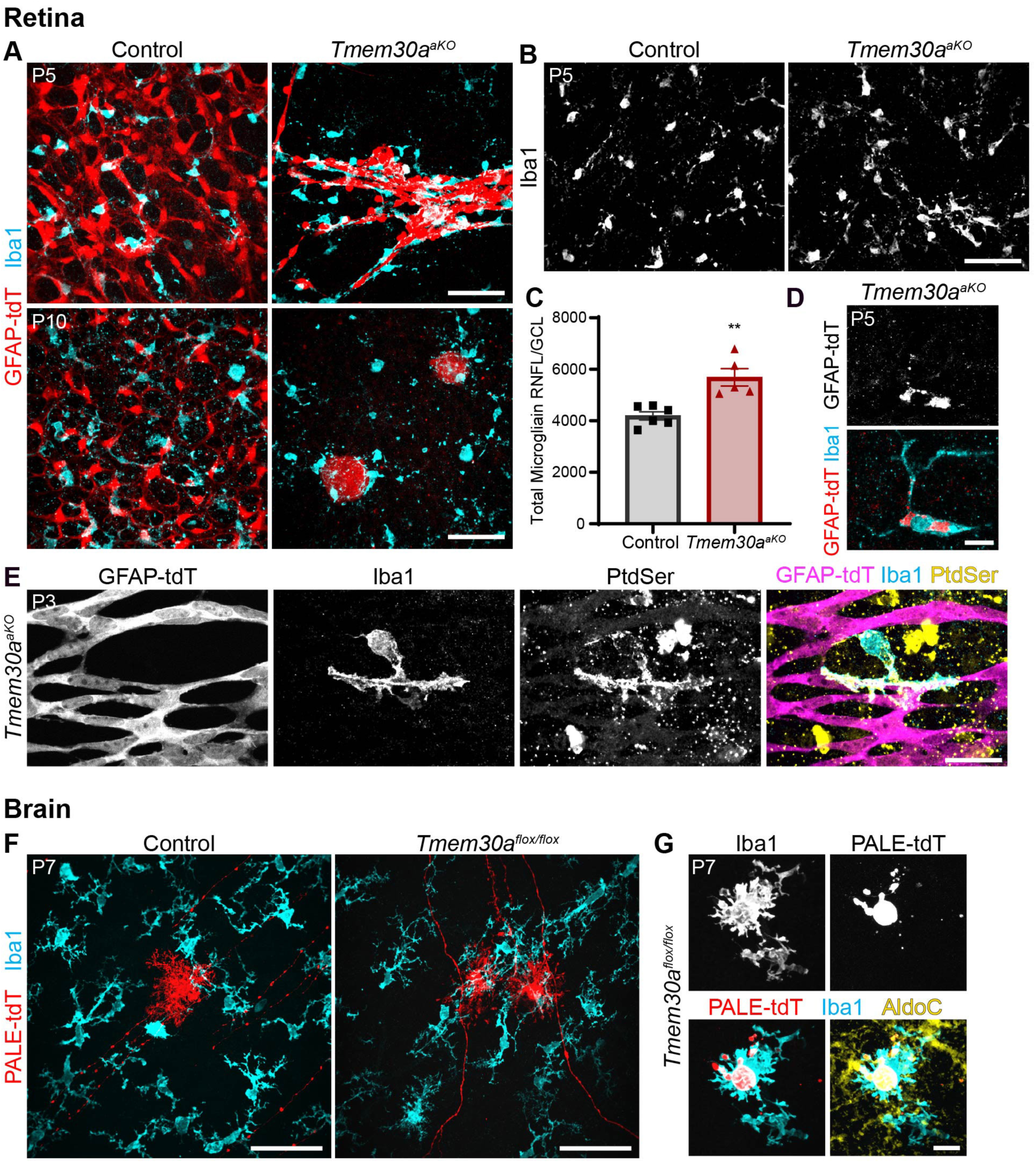
Microglia are recruited to PtdSer-expressing astrocytes. A) Microglia interact preferentially with astrocyte aggregates in *Tmem30a^aKO^* retinas. Left, representative images showing uniform distribution of RNFL microglia (Iba1, cyan) and astrocytes (GFAP-tdT, red) in control retina at P5 (top) and P10 (bottom). In *Tmem30a^aKO^* retinal regions containing astrocyte aggregates (right panels), microglia are no longer spaced uniformly but are instead selectively localized to the astrocyte clumps. B,C) RNFL microglia numbers are increased in *Tmem30a^aKO^* mutants. B: Representative images of Iba1+ microglia in control and *Tmem30a^aKO^*retinas at P5. Images show maximum-intensity projections of Z-stacks through RNFL and GCL (∼10-13 µm). C: Quantification of total microglia number from images similar to B, estimated using densities weighted by eccentricity. Statistics: two-tailed *t* test, \*\**p* = 0.002. D) Microglia engulf astrocyte material in*Tmem30a^aKO^* retina. Example image of a microglial cell (Iba1, cyan) containing large chunks of astrocyte debris (GFAP-tdT, red). E) Microglia contact astrocytes at sites of PtdSer exposure. Example image from a P3 *Tmem30a^aKO^* retinal wholemount that was stained live with anti-PtdSer (yellow), then fixed and processed for microglia (Iba1, cyan) and astrocyte (GFAP-tdT, magenta) markers. A microglial cell makes extensive contact along an astrocytic process; PtdSer localizes selectively to the interface between the two cells. F) Brain microglia also interact preferentially with *Tmem30a* mutant astrocytes. Example images from P7 *Ai14* control and *Tmem30a^flox/flox^* PALE brains electroporated with Cre. In controls, microglia (Iba1, cyan) are distributed uniformly despite presence of astrocytes expressing PALE-tdT reporter (red). In *Tmem30a^flox/flox^* mice, microglia accumulate in the vicinity of transfected mutant astrocytes. G) Brain microglia engulf *Tmem30a* mutant astrocytes. Example image shows a microglial cell (Iba1, cyan) in the cortex of a P7 *Tmem30a^flox/flox^*PALE brain, containing large chunks of Cre+ astrocyte debris (PALE-tdT, red; AldoC, pan-astrocyte marker, yellow). The debris inclusions found in microglia in *Tmem30a^flox/flox^* PALE mice were much larger than those found in control PALE mice. Scale bars: A,B,F: 50 µm; D,G: 10 µm; E: 25 µm.

Importantly, microglial aggregation behavior was not limited to the retina: Microglia in the cortex of Cre-electroporated *Tmem30a^flox/flox^* animals also accumulated in the vicinity of *Tmem30a* knockout astrocytes (Figure 6F). Additionally, we found microglia with large inclusions of PALE-tdT+ debris, the scale of which was not seen in control animals (Figure 6G). Together, this evidence strongly supports the view that forced PtdSer exposure leads to death of both retinal and cortical astrocytes through a mechanism involving recruitment and phagocytic activity of microglia.

### PtdSer binding protein MFGE8 is required for PtdSer-mediated astrocyte death

To study the molecular mechanism by which microglia eliminate astrocytes, we investigated the known molecular pathways by which phagocytes recognize and engulf PtdSer+ cells. Microglia express numerous different receptors that can mediate this process (Brown & Neher, 2014; Lemke, 2013). These phagocytic receptors do not recognize PtdSer directly; rather, cells that have exposed cell-surface PtdSer become opsonized by soluble PtdSer-binding proteins that serve as ligands for the microglial receptors. Because there are several different opsonins and several different receptors, the system is highly molecularly redundant. We initially investigated the TAM-family tyrosine kinase receptor MERTK, which recognizes the PtdSer-binding opsonins Gas6 and Protein-S, but we found that *Mertk* mutant mice lack retinal astrocyte phenotypes (Puñal et al., 2019). We therefore turned to a second pathway, mediated by the PtdSer-binding opsonin MFGE8 and its microglial integrin receptors (Lemke, 2013). We considered MFGE8 a good candidate because *Mfge8* RNA and protein are expressed by several different cell types within the RNFL, including pericytes, vasculature, microglia, and astrocytes themselves (Li et al., 2024; Motegi et al., 2011). However, *Mfge8* null mutant mice also lacked any defects in astrocyte death (Supplemental Figure 5A,B). To test for redundancy between these pathways, we generated *Mfge8; Mertk* double mutant mice. Again, no astrocyte death phenotypes were observed (Supplemental Figure 5C,D). This was perhaps not surprising, because there are at least two other opsonin-receptor pathways by which microglia could potentially recognize and eliminate PtdSer+ astrocytes even when the TAM and MFGE8 pathways are both impaired (Anderson et al., 2019; Gnanaguru et al., 2023; Nagata, 2018; Païdassi et al., 2008; Schafer et al., 2012; Scott-Hewitt et al., 2020; VanRyzin et al., 2019).

Given this redundancy and the impracticality of generating quadruple mutant mice, we sought an alternate strategy to determine the mechanism by which microglia kill PtdSer+ astrocytes. To this end, we used *Tmem30a^aKO^* mice as a sensitized background to probe the effects of blocking individual PtdSer signaling pathways. Loss of a single pathway in the wildtype context may not degrade PtdSer recognition enough to impair astrocyte removal. But in the context of forced PtdSer exposure, where demands on the PtdSer recognition mechanism are higher, it is plausible that loss of a single PtdSer recognition pathway could suffice to at least partially block microglia from causing astrocyte death. To test this idea, we generated a *Tmem30a^aKO^*; Mfge8^−/–^double knockout line. Strikingly, astrocyte death was substantially rescued in the double knockout animals, despite the continued forced exposure of cell-surface PtdSer. While GFAP retinal coverage did not fully return to control levels in double mutants, indicating that there is still a role for other PtdSer recognition pathways, the astrocyte network was largely intact and covered far more of the double mutant retina than in *Tmem30a^aKO^* single mutants (Figure 7A-C). These observations indicate that microglia recognize PtdSer on astrocyte cell surfaces using the MFGE8 pathway, and that activation of this pathway is needed for PtdSer-mediated death.

**Figure 7:**
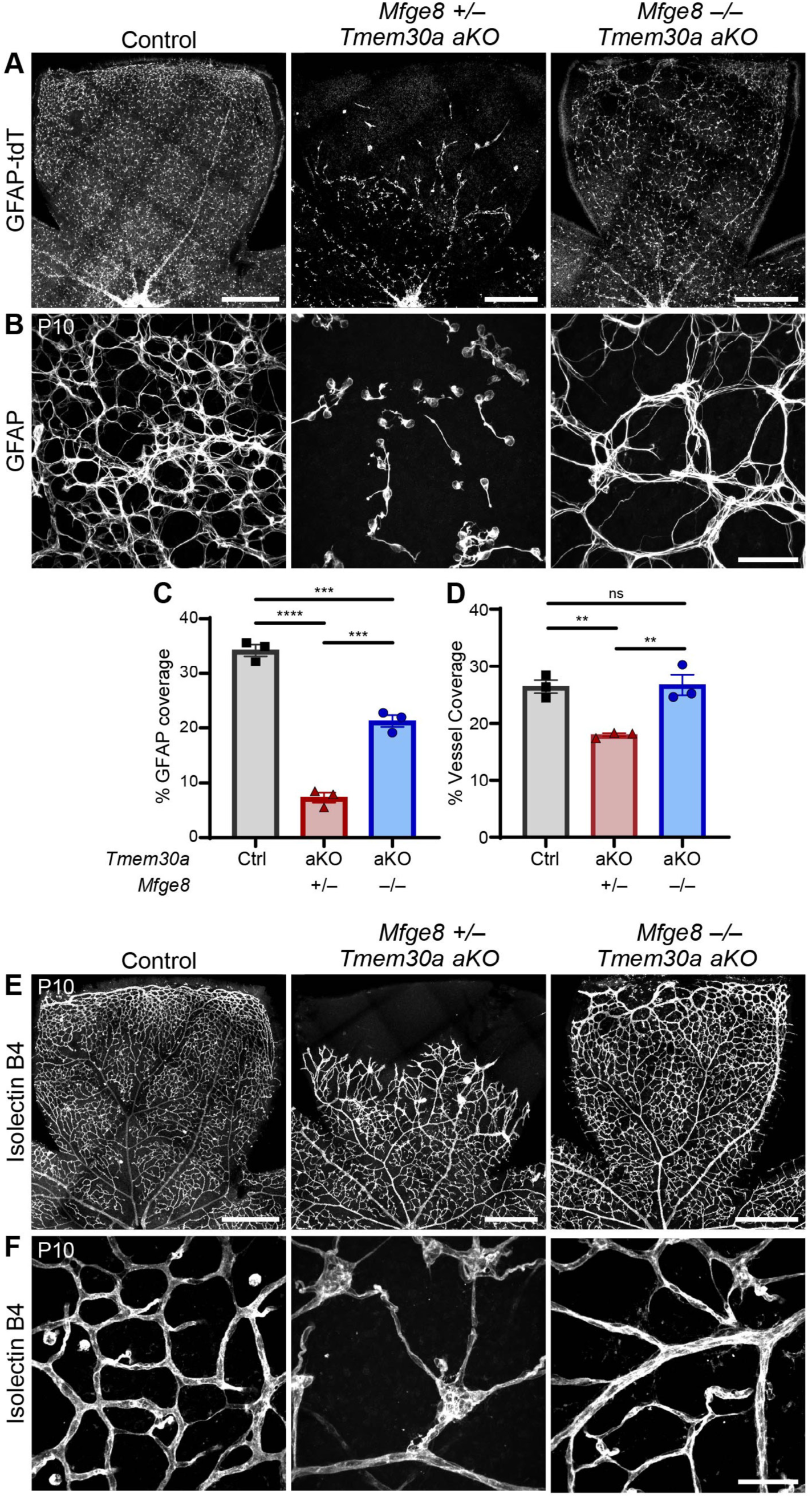
PtdSer binding protein MFGE8 is required for PtdSer-mediated astrocyte death. A,B) Example images of RNFL showing restoration of astrocyte coverage when *Mfge8* is removed from *Tmem30a^aKO^* mutants. A, low power overviews using GFAP-tdT to mark astrocytes; B, high magnification images showing GFAP immunostaining, representative of those used for coverage quantification (C). P10 control retina (left) is fully occupied by astrocytes (A) which make a dense mesh network (B). *Mfge8^+/–^*; *Tmem30a^aKO^* mice (center) phenocopy *Tmem30a^aKO^* single knockouts (Fig. 2C,D; Fig. 5A,B) in their loss of astrocyte coverage. *Mfge8^−/–^*; *Tmem30a^aKO^*double knockouts (right) exhibit a partial but substantial rescue of astrocyte occupancy (A) and mesh network coverage (B). C,D) Quantification of GFAP coverage (C) or vessel coverage (D) across the three indicated genotypes. Graphs show average coverage from a series of images similar to B or F, collected to sample uniformly across the retina. GFAP (C) and vascular (D) coverage were both greatly diminished in *Tmem30a^aKO^* mutants that retained a functional copy of *Mfge8* (control vs. *Mfge8^+/–^*; *Tmem30a^aKO^*, GFAP \*\*\*\**p* = 3.7 x 10^−6^, vessels \*\**p* = 0.007), confirming results of Fig. 5B,D. Complete removal of *Mfge8* on the *Tmem30a^aKO^* background (double mutants, blue) significantly improved both astrocyte and vessel coverage compared to *Mfge8* heterozygotes on this background (red; GFAP \*\*\**p* = 1.79 x 10^−4^, vessels \*\**p* = 0.006). GFAP coverage in double mutants was not restored all the way to control levels (control vs. double mutant, \*\*\**p* = 2.85 x 10^−4^), but vessel coverage was fully rescued (control vs. double mutant, *p* = 0.985). Statistics: one-way ANOVA (GFAP \*\*\*\**p* = 5.2 x 10^−6^, vessels \*\**p* = 0.004) followed by Tukey’s post-hoc tests corrected for multiple comparisons, *p*-values noted above. E,F) Example images showing rescue of RNFL vascular phenotypes when *Mfge8* is removed from *Tmem30a^aKO^* mutants. E, low power overviews of vessel (Isolectin B4) coverage. F, high magnification images representative of those used for vessel coverage quantification (D). In P10 control retinas (left), vessels are fully invested throughout the RNFL. *Mfge8^+/–^*; *Tmem30a^aKO^*mice (center) phenocopy *Tmem30a^aKO^* single knockouts (Fig. 5C-F) in their loss of vessel coverage and formation of hypertrophic tufts. Double mutants (right) are largely restored to control state, aside from some small regions at the far retinal margin. Because of their size and marginal location, these unrescued regions were not well captured in our quantitative analysis (D). Error bars: mean ± SEM. Scale bars: A,E: 500 µm; B,F: 50 µm.

We considered the possibility that the rescue of GFAP+ astrocytic material in double mutant mice reflects a requirement for MFGE8 in the microglial clearance of dead astrocytic debris, rather than a requirement in the death mechanism itself. To distinguish between these possibilities, we investigated whether the rescued astrocytes are functionally normal. If deletion of MFGE8 is truly preventing astrocytes from dying, the rescued astrocytes should be able to generate a functional template for retinal angiogenesis. Indeed, analysis of vascular anatomy revealed that, in striking contrast to *Tmem30a^aKO^*single mutants, vessel coverage in double mutants was nearly indistinguishable from controls (Figure 7D-F). The only minor appreciable differences between double mutant and control vasculature were slight disruptions at the far retinal periphery (Figure 7E). Thus, while astrocyte coverage was not rescued fully to control levels, these results clearly demonstrate that the rescued astrocytes are able to fully support angiogenesis. Importantly, we did not observe any hypertrophic tufts in double mutants, indicating a rescue of disease-like pathological features as well (Figure 7D-F). Given their ability to mediate normal vascular development, we conclude that astrocytes are indeed rescued from PtdSer-mediated death when MFGE8 is absent. Altogether, these findings establish MFGE8 as a key part of the molecular pathway by which microglia kill developing astrocytes upon exposure of cell-surface PtdSer.

## DISCUSSION

Developmental death of retinal astrocytes occurs by an unusual non-apoptotic mechanism. Here we have identified PtdSer as a cell-surface “kill-me” signal that serves as a central initiator of the “death by phagocyte” pathway responsible for culling astrocytes. This mechanism is used not only in neonatal retina, but also by astrocytes of the developing cerebral cortex, suggesting that it is a broad and general mechanism for astrocyte death. Furthermore, astrocytes appear uniquely sensitive to this death mechanism, because forced PtdSer exposure in neurons does not cause cell death – either in the retina (Figure 3A,B) or in the brain (Li et al., 2021; Park et al., 2021). Our anatomical and genetic evidence support the following model of astrocyte death: During the developmental death window, a subset of astrocytes exposes cell-surface PtdSer, which leads these astrocytes to become opsonized by the PtdSer-binding protein MFGE8. Opsonization, in turn, recruits microglia to engulf and kill the PtdSer-exposing astrocytes. This PtdSer-MFGE8 pathway is responsible for astrocyte death, rather than simply for clearance of dead cells, as shown by the substantial increase in surviving, angiogenesis-competent astrocytes in *Mfge8; Tmem30a^aKO^* double mutants. Because astrocyte survival and morphology were not fully rescued by the removal of MFGE8, it is likely that other PtdSer-binding opsonins, such as Gas6, Protein S, and C1q, are also involved (Brown & Neher, 2014; Gaboriaud et al., 2011; Lemke, 2019; Nagata, 2018).

In the model delineated above, PtdSer externalization is the event that initiates astrocyte death because it creates the initial signal for microglial recruitment. Consistent with this initiation idea, experimentally increasing the fraction of PtdSer+ astrocytes accelerates death. However, the number of astrocytes that ultimately die remains unchanged in *Tmem30a^aKO^* mutants relative to wildtype mice, despite continued PtdSer exposure. This cessation of death has two important implications. First, it demonstrates that *Tmem30a* deletion does not kill astrocytes through a non-specific mechanism involving cellular dysfunction or degeneration. Second, it shows that MFGE8-dependent microglial elimination of PtdSer+ astrocytes occurs only under specific conditions: When astrocyte numbers fall beyond a certain point, the mechanism can be suppressed to prevent over-elimination. Thus, PtdSer exposure is best viewed not as a guarantee of death, but rather as a pro-death signal whose outcome depends on context. Further support for this view comes from our flow cytometry assay: The fraction of PtdSer+ astrocytes at P5-P6 was ∼10%, far higher than the number of astrocytes that are dying at any given time (Puñal et al., 2019), which suggests that many astrocytes transiently externalize PtdSer without dying. The probability of progressing from externalization to death likely depends on factors including: 1) the duration of PtdSer exposure (Segawa & Nagata, 2015); 2) the number and/or phagocytic transcriptional state of local microglia (Martineau et al., 2026); and 3) the presence of competing “don’t eat me” signals (Jiang et al., 2022).

The finding that PtdSer exposure promotes astrocyte death raises the question of how astrocytes choose to engage in PtdSer externalization. Prior studies have shown that the probability and duration of PtdSer exposure depends on the balance between flippase and scramblase activity (Figure 1B) (Segawa & Nagata, 2015). Therefore, one reasonable possibility is that live astrocytes have higher baseline scramblase activity, and/or lower baseline flippase activity, than do other cell types that lack the propensity to externalize PtdSer. In this scenario, PtdSer exposure and the decision to die would be probabilistic: all astrocytes would have a similar propensity to occasionally place PtdSer on the cell surface for a brief period of time, with the time constants for exposure and internalization determined by the two enzymatic activities. The cells that happened to leave PtdSer on the surface longest would be most vulnerable to detection and removal by microglia. However, we do not rule out the existence of other types of signals capable of forcing longer-term PtdSer exposure in a subset of astrocytes, thereby marking them specifically for removal. The extent to which death occurs stochastically, or in a targeted manner, remains to be clarified.

Our studies of vessel development demonstrate a critical functional role for proper regulation of astrocyte PtdSer exposure: In *Tmem30a^aKO^*mice, formation of the primary vascular plexus is not only delayed, but the resulting vessel network is also mispatterned. These effects on vessels likely arise at least in part from the change in astrocyte death period kinetics that occurs in *Tmem30a^aKO^* mice. During RNFL angiogenesis, astrocytes are the key source of the pro-angiogenic factor VEGF-A, without which vessel development cannot proceed normally (Paisley & Kay, 2021; Rattner et al., 2019; Stone et al., 1995). The timing of the astrocyte death window suggests that a key function of death is to diminish VEGF-A signaling once primary angiogenesis is complete. Accelerated death would therefore be expected to hasten the drop in VEGF-A expression, thereby depriving the developing vasculature of this critical signal at a time when it is still needed. An additional consequence of forced PtdSer exposure was evident from the fact that, even in regions of *Tmem30a^aKO^* retina where astrocytes are present, vessels still were not patterned normally. This vascular defect likely reflects the altered morphology of *Tmem30a^aKO^*mutant astrocytes, particularly the loss of their fine arbors that normally form a template for vessel growth (Fruttiger et al., 1996; Gerhardt et al., 2003; O’Sullivan et al., 2017). We conclude from this loss of fine arbors, together with our observations of microglia wrapping PtdSer+ astrocyte arbors (Figure 6E), that PtdSer exposure likely drives astrocyte arbor pruning in addition to soma removal. Arbor pruning may occur in parallel to death, or it may instead be a death precursor that needs to happen first before microglia can ingest the astrocyte soma. Either way, the vascular phenotypes in *Tmem30a^aKO^* retina indicate that arbor and soma removal are both functionally important.

As an example of this functional importance, vascular anomalies resembling ROP pathologies were observed at the angiogenic wavefront of *Tmem30a^aKO^* mice. One such anomaly was the retardation of the angiogenic wavefront, leaving a large portion of the peripheral retina without vessels – a finding that is common in clinical images from preterm infants with ROP. A second anomaly observed both in mutant mice and ROP clinical imaging was the presence of intraretinal neovascularization at the angiogenic wavefront (Figure 5) (Chen et al., 2024). In mice, these neovascularizations usually lacked underlying astrocytes, presumably because the astrocytes died soon after vessels grew over them. Thus, our results imply that loss of astrocyte support can induce nascent vessels to undergo pathological changes that contribute to ROP phenotypes. In addition to loss of support, an excess of astrocytes can also be deleterious to vessels in ways that resemble different features of ROP (Duan & Fong, 2019; Duan et al., 2014; Fruttiger et al., 1996; Hellström et al., 2013; McMenamin et al., 2016; Morita et al., 2016; Perelli et al., 2021; Stone et al., 1995; West et al., 2005). Thus, our findings underscore the importance and disease relevance of astrocyte-vascular communication not only during initial vessel outgrowth, but also during vessel differentiation.

As a mechanism that can initiate death, PtdSer-MFGE8 signaling is well-positioned to serve as a cue for the opening and progression of the astrocyte death window. However, open questions remain as to how the final number of astrocytes is determined, and how the death period closes once this number is reached. Non-mammalian vertebrate species can make do without any retinal astrocytes at all (Schnitzer, 1988; Todd et al., 2016), so the fact that some astrocytes survive the death period suggests that they must possess important functions in the adult mammalian retina (Holden et al., 2023). There are at least three plausible mechanisms that might close the death period or otherwise serve to prevent excess astrocyte elimination. First, contact with vasculature could provide a trophic cue, as was previously shown for cortical astrocytes (Foo et al., 2011). Supporting this idea, we noted that, in some *Tmem30a^aKO^*retinas, astrocytes lining the major vessels in central retina appeared to be selectively preserved (Supplemental Figure 2H). Second, developmental changes within the microglia population might make them less responsive to PtdSer, MFGE8, and/or other astrocyte-derived cues, thereby reducing their ability to kill. This is plausible because developing microglia express distinctive pro-phagocytic gene profiles that are not present in adulthood (Hammond et al., 2019; Li et al., 2019; Martineau et al., 2026). Third, developing retina exhibits an increase in “don’t eat me” signaling during the second postnatal week, mediated through the SIRPα-CD47 pathway, which could inhibit microglia killing activity (Jiang et al., 2022). Future work will be needed to learn the extent to which these or other mechanisms collaborate with PtdSer signaling to dictate the extent of astrocyte death, as well as to assess the impact of PtdSer-mediated death on development of astrocytes and vasculature within other CNS regions.

## METHODS

### Experimental Subjects

#### Animals

The Institutional Animal Care and Use Committee at Duke University reviewed and approved all experimental procedures involving animals. Mice of both sexes were used for experiments. We did not include or exclude animals on the basis of sex. Mice were housed on a 12-hour light/dark cycle with unrestricted access to food and water. All mouse strains were propagated by backcrossing to C57Bl/6J, unless otherwise specified. Upon arrival in our colony, each strain underwent genotyping for *rd1* and *rd8* retinal degeneration mutations. If present, these mutant alleles were eliminated through selective breeding.

For Cre-dependent expression of the tdTomato fluorescent reporter, we obtained *Rosa26^Ai14^* mice (Madisen et al., 2010) from the Jackson Labs repository. GFAP-Cre (Tg(GFAP-cre)25Mes) (Lee et al., 2008), also from Jackson Labs, was the astrocyte-specific Cre line used in these studies. *Tmem30a^tm1a(KOMP)Wtsi^*mice (INFRAFRONTIER Consortium, 2015) were obtained from The European Mouse Mutant Archive (EMMA) Repository and were used to generate *Tmem30a^aKO^*mutants in the following manner. First, they were crossed to ACTB:Flpe (Rodríguez et al., 2000) mice to remove FRT sites, thereby generating a *Tmem30a^flox^* allele; and to Actin-Cre (Lewandoski et al., 1997) mice, thereby generating a *Tmem30a^null^* allele. Subsequently, to produce breeders that could be used for the generation of experimental mice, *Tmem30a^flox^* animals were crossed to Ai14 and to GFAP-Cre. Because this Cre strain occasionally exhibits unwanted recombination activity in the germline, care was taken to use only first-generation progeny of *Tmem30a^flox^* x GFAP-Cre crosses as breeders for experimental matings. Additionally, to reduce the possibility of germline recombination, breeders for experimental matings were sometimes generated by crossing *Tmem30a^null/+^*mice with GFAP-Cre, to produce GFAP-Cre; *Tmem30a^+/null^* breeders. Finally, *Tmem30a^aKO^* mutants and littermate controls were generated by crossing *Tmem30a^flox^*; *Rosa26^Ai14/Ai14^*animals with either GFAP-Cre; *Tmem30a^flox/+^*, or GFAP-Cre; *Tmem30a^null/+^* mice. Thus, the genotype of *Tmem30a^aKO^*mutants was either GFAP-Cre; *Tmem30a^flox/–^* or GFAP-Cre; *Tmem30a^flox/flox^*. Control genotypes included GFAP-Cre; *Tmem30a^+/+^* and GFAP-Cre; *Tmem30a* heterozygotes, which showed no cell death phenotype (Supplemental Figure 2E). All experimental animals carried *Rosa26^Ai14^*to track Cre expression in astrocytes. During timepoints examined in this study, *Tmem30a^aKO^* animals were behaviorally and physically normal. Around 3 weeks of age, *Tmem30a^aKO^* mice started to look unhealthy and survival dropped off by 1 month of age, suggesting that, in addition to the developmental functions reported here, the gene is also required in mature astrocytes or other non-CNS cell types expressing the GFAP-Cre transgene.

To generate cholinergic neuron-specific *Tmem30a* mutants (*Tmem30a^sacKO^*) or OPC-specific *Tmem30a^opcKO^* mutants, *Chat^Cre^* (*Chat^tm2(cre)Lowl^*) (Rossi et al., 2011) or *Olig2^Cre^* (*Olig2^tm1.1(cre)Wdr^*) (Zawadzka et al., 2010) animals were used in place of GFAP-Cre animals in the breeding strategy described above. Similar to *Tmem30a^aKO^*, we noted a deterioration in the health of *Tmem30a^sacKO^* animals during weeks 2-3, whereas *Tmem30a^opcKO^* mice remained apparently healthy during our experimental timeframe.

*Mfge8^Gt(KST227)Byg^* (Atabai et al., 2005; Nandrot et al., 2007) mice were rederived from frozen sperm at the Mouse Breeding Core at Duke University on a B6SJLF1/J hybrid background. After derivation, at least 5 backcrosses to C57Bl6/J were performed to establish the line on a pure C57Bl6/J background which was confirmed by SNP genotyping at the DartMouse facility (Trask et al., 2010) This backcrossed *Mfge8* strain was then bred with *Tmem30a^flox^*mice, generating the *Mfge8; Tmem30a^flox^* double mutant line; and with the *Mertk^tm1Grl^* knockout strain (Lu et al., 1999), obtained from Jackson Labs, to make *Mfge8;Mertk* double mutant animals.

#### Human OCT imaging data

OCT and OCTA images shown in this study were obtained through re-analysis of an imaging dataset that was reported previously (Chen et al., 2024). No new human subjects were enrolled for this study. The study was approved by the Duke Health Institutional Review Board and study participants were imaged with informed consent from parents/guardian. The study adhered to the Health Insurance Portability and Accountability Act and all tenets of the Declaration of Helsinki.

### Method Details

#### Postnatal Astrocyte Labeling by Electroporation (PALE)

Late P0/early P1 *Tmem30a^flox/+^*; *Rosa26^Ai14/+^* and *Tmem30a^flox/flox^*; *Rosa26^Ai14/+^* littermate mouse pups were sedated using ice anesthesia. 1 µL of pCag-Cre plasmid DNA (gift from Connie Cepko, Addgene plasmid #13775) mixed with Fast Green dye was injected into the lateral ventricle of one hemisphere of the brain using a pulled glass pipette. Following DNA injection, electrodes were placed on the head so that the positive terminal was above the visual cortex and the negative terminal was below the chin of the animal. 5 discrete 50 msec pulses of 100V spaced 950 msec apart were applied. Pups were placed on a heating pad to recover, then returned to their home cage and monitored until collection at P7 or P14. All animals that appeared healthy at the time of collection were processed for analysis. Prior to downstream processing, brains were examined for the presence of electroporated cells (PALE-tdT+ cells) using a Leica M165 FC epifluorescence dissecting microscope. Brains with no labeled astrocytes were excluded from the study.

#### Histology

For all retina histology experiments, P1-P6 neonatal mice were euthanized by ice anesthesia followed by decapitation and immediate enucleation. Mice P7 and older were euthanized by isoflurane anesthesia followed by decapitation and immediate enucleation. Whole eyes were fixed in 4% paraformaldehyde (PFA) in 1X PBS on ice for 90 minutes, then washed at least 2 times in 1X PBS. Once fixed, eyes were either processed immediately for immunohistochemistry, or they were stored at 4°C in 0.02% sodium azide in 1X PBS until processing. For anti-PtdSer labeling, the same euthanasia protocols were followed, but after enucleation, retinas were immediately dissected out of the eye cup in oxygenated Ames’ medium (with sodium bicarbonate added) and were fixed only after incubation with the anti-PtdSer antibody (see below for details).

After fixation, eyes were either processed for whole-mount immunohistochemistry, or for cryosectioning followed by immunostaining on slides. For retinal sections, the cornea was removed and the lens and vitreous extracted from the eyecup. Eyecups were cryoprotected in 30% sucrose for at least 2 hours and then frozen in Tissue Freezing Medium. Retinas were cryosectioned at 20 µm and mounted on Superfrost Plus slides. Whole-mount retinas were prepared for downstream staining by removing the cornea, extracting the lens and vitreous, and detaching the retina from the eyecup.

For brain histology, mice were anesthetized with isoflurane before being transcardially perfused with 1X PBS followed by 4% PFA in 1X PBS. After perfusion, mice were decapitated and whole brains were removed. Brains were postfixed overnight in 4% PFA in 1X PBS at 4°C. After postfixation, brains were washed 3-5 times (30-60 minutes each) in 1X PBS. Brains were either 1) embedded in 2% agarose and 80 µm coronal sections were collected on a Leica VT 1200 S vibratome; or 2) cryoprotected in 30% sucrose overnight, frozen in Tissue Freezing Medium, and 80 µm coronal sections were collected on a Leica CM1950 cryostat. The sequential order of the brain sections was preserved. For electroporated brains, sections were chosen for analysis by determining the beginning and end of the electroporated patch in the cortex using a fluorescence dissection microscope and then choosing 3 sections that were evenly spaced within the area of the patch.

#### Immunohistochemistry

For immunohistochemistry, retinal tissue was blocked with a solution of 3% normal donkey serum, 0.3% Triton-X 100, and 0.02% sodium azide in 1X PBS at room temperature. Blocking occurred for 2 hours for whole-mount retinas and 30 minutes for retinal sections. After blocking, retinal tissue was incubated with primary antibodies diluted in blocking solution (see Key Resources Table for dilutions) either for 5-7 days for wholemount retinas or overnight for retinal sections. Primary incubation occurred at 4°C and included rocking for wholemount retinas. Retinal tissue was washed after antibody incubation at least 3 times in 1X PBS. Secondary antibodies and Hoechst 33258 were diluted in a solution of 0.3% Triton-X 100 in 1X PBS. All secondary antibodies and Hoechst 33258 were used at a 1:1000 dilution. Secondary antibodies were applied either overnight at 4°C for wholemount retinas or for 2 hours at room temperature for retinal sections. Finally, tissue was washed at least 3 times at room temperature before mounting tissue for image acquisition.

For anti-PtdSer labeling, live retinas were blocked with 4% BSA and rat anti-mouse Fc block (1:400) in oxygenated Ames’ solution (with sodium bicarbonate added) for 15 minutes at room temperature. Then, retinas were incubated with anti-PtdSer diluted in blocking solution for 20 minutes at room temperature, as previously described (Ruggiero et al., 2012). After incubation with the primary antibody, retinas were quickly washed 5 times with oxygenated Ames’ to remove excess antibody, then fixed in 4% PFA in 1X PBS for 45 minutes on ice. After fixation, retinas were washed 3 times in 1X PBS for 5 minutes each. Immunohistochemistry was performed as described above, with the addition of other primary and secondary antibodies as needed, and using anti-mouse secondary to detect labeling with the anti-PtdSer primary antibody. In anti-PtdSer labeled retinas, astrocytes were analyzed only outside of the vessel network to avoid overlapping PtdSer+ vessel signal. As a positive control, apoptotic cells were identified by their pyknotic morphology visualized with Hoechst 33258 nuclear labeling. Apoptotic cells were routinely labeled by anti-PtdSer, as expected (Figure 1E).

#### Image acquisition and processing

Following immunohistochemistry, four radial cuts separated by 90 degrees were made in whole retinas to allow for flattening of the retina (Figure 1B). Cut retinas were placed astrocyte-side up on nitrocellulose filter paper and paintbrushes were used to carefully lay each lobe flat. Retinas were then mounted on a slide and coverslipped using Fluoromount-G mounting media. The mounting media was allowed to cure for at least 2 hours before the edges of the coverslip were nail polished to hold the coverslip in place. Nikon A1, A1R, or Olympus FV300 confocal microscopes were used to image retina sections and wholemounted retinas. Sections and wholemounts were imaged with a 60x oil objective and Z-stacks were collected at a Z-resolution of 0.3–1µm.

Whole retina tilescans (e.g. Figures 2A, 5E, 7A,D), or tilescans of whole brain sections, were acquired with a 20x air objective and stitched together into a single image using software (Nikon Elements) provided by the microscope manufacturer. Images were imported to Fiji (Schindelin et al., 2012) for processing and analysis. Images selected for publication are maximum-intensity projections of a subset of the Z-slices within the confocal stack, with the minimum number of Z-planes included to show the structure being discussed, unless otherwise noted. After Z-projection, images were de-noised by median-filtering (0.5–2.0-pixel radius). Composite images were produced and minor adjustments to brightness and contract were made in Fiji software. For analysis and quantification, original stacks were used instead of Z-projections, unless otherwise noted.

#### Analysis of human OCTA images

The infant was imaged at bedside in the nursery with an investigational 200kHz swept-source handheld OCT system (UC3, Duke University, Durham, NC). OCTA images were processed and generated using custom MATLAB software, and cross-sectional B-scans were segmented automatically (using BabyDOCTRAP, Duke University, Durham, NC) followed by manual correction by experienced grader as previously reported (Chen et al., 2024).

#### Flow cytometry

For flow cytometry analysis, P5–P6 live retinas were removed from eyecups and dissociated using papain incubation and trituration, and live-stained with conjugated antibodies recognizing cell-surface antigens. Antibodies to PDGFRα were used to label astrocytes, and antibodies to Thy1 were used to label retinal ganglion cells as a positive control for Annexin V labeling (see below). Cells were also stained with a live/dead staining kit (Invitrogen Cat# L34965). All manipulations of retinas and retinal cells were performed in solutions made with oxygenated Ames’ media (with sodium bicarbonate added), except for Annexin V-APC labeling, which was performed in Annexin V Binding Buffer.

After staining, dissociated cells were analyzed using a BD FACSAria flow cytometer. Data was imported into FlowJo software for further offline analysis. First, a compensation matrix was created and applied to all data from each experiment. Then, dead cells were removed from analysis based on live/dead staining. Gates were determined based on innate fluorescence levels of unstained cells for live/dead and PDGFRα channels. During this analysis, we noticed that *Tmem30a^aKO^*astrocytes had higher PDGFRα staining than their control littermates. This is potentially due to the reduced number of vessels in these knockout animals. Incoming vessels are known to provide immature astrocytes with multiple maturation cues (both oxygen- and contact-mediated (Paisley & Kay, 2021)), so without these signals, it is possible that astrocytes are not maturing, thus not downregulating *Pdgfra* expression. Because of this, PDGFRα signal from control astrocytes was used to draw the astrocyte gate.

To determine where the boundary between Annexin V+ and Annexin V-astrocytes should be drawn, retinal ganglion cells stained with Thy1 and Annexin V were analyzed within the same experiment as a positive control. Retinal ganglion cells are induced to undergo apoptosis as a result of the dissociation process, due to the severing of their axons. Thus, ganglion cells are a suitable positive control for Annexin V staining. We observed a clear separation between Annexin V+ and Annexin V-retinal ganglion cells (Supplemental Figure 1D), allowing us to determine where the Annexin V gate should be drawn for the astrocyte population based on an objective independent criterion. Many Annexin V+ ganglion cells were negative for the dead cell fluorescent stain, consistent with the externalization of PtdSer at an early stage of apoptosis before the loss of membrane integrity.

Histograms of Annexin V-APC fluorescence intensity for PDGFRα+ cells were used to determine the percentages of Annexin V+ and Annexin V-cells in the astrocyte population. For more information about the gating strategies used, see Supplemental Figure 1.

#### Expression of flippase genes

Expression of genes encoding P4-ATPase family members were assessed using the Mouse Retinal Cell Atlas dataset (Li et al., 2024), accessed through the Broad Single Cell Portal, which was also used to generate the dot plot graph.

### Quantification and Statistical Analyses

#### Statistics

Error bars are expressed as mean ± SEM. For all analyses, α = 0.05. Sample sizes are reported either by individual points on graphs or in figure legends. Statistical tests, including post-hoc analyses, and *p*-values are reported in every figure legend. All *t* tests were two-tailed. One-way and two-way ANOVAs without matching were used, followed by appropriate post-hoc tests if ANOVAs showed significant main effects or interactions. Asterisks denote statistical significance (\**p* ≤ 0.05; \*\**p* ≤ 0.01; \*\*\**p* ≤ 0.001; \*\*\*\**p* ≤ 0.0001). Data were analyzed using GraphPad Prism software.

#### Quantification of retinal astrocyte density and total astrocyte number

For retinal astrocyte counts, we sampled RNFL astrocyte density from three distinct regions: central, middle, and peripheral retina. Confocal Z stacks, typically 3-4 per region, were acquired using a 60x objective on the Nikon confocal system, generating a ROI that was 212.13 x 212.13 µm. Central images were chosen from retina tissue proximal to the optic nerve head (point where blood vessels enter the retina). Regions containing major blood vessels were avoided because they could cause local distortions in astrocyte distribution. Middle images were acquired at the approximate midpoint between the optic nerve head and the peripheral edge of each lobe. Peripheral images were selected approximately 2–3 astrocyte cell body lengths away from the edge of the retina. In rare circumstances, only 2 images were available in a given region. Only undamaged areas of the retina were analyzed. Astrocyte density was quantified by hand for each image using Fiji software to manually mark each cell.

The density measurements from center, middle, and peripheral regions were used to estimate the total number of astrocytes in each retina, using the approach we have previously described (O’Sullivan et al., 2017; Puñal et al., 2019). First, the average density per region was calculated for the 3-4 images from that region. Next, the average densities per region were used to compute an overall weighted average density for each retina. The weighting takes into account the fact that the concentric rings from which the center, middle, and peripheral images were sampled are not of equal size. The weighting percentages, calculated as described previously, were as follows: central = 11%, middle = 33%, and peripheral = 56% (O’Sullivan et al., 2017; Puñal et al., 2019). This weighted average density was multiplied by the total area of the retina to estimate the total number of astrocytes. Retina areas were measured in Fiji by drawing perimeters with the Freehand or Polygon selection tools on tilescan images of wholemount retinas. Similar methods were followed for the quantification of total microglia number.

#### Quantification of colocalization and coexpression of astrocyte markers

For analysis of Sox9 and GFAP-tdT coexpression, image stacks were acquired as described above. Astrocytes labeled by Sox9 were counted first. Cells labeled by GFAP-tdT were counted independently of Sox9 labeling. Colocalization fractions were determined based on the percentage of cells that were determined to be both Sox9+ and GFAP-tdT+, without weighting based on eccentricity (as in the total cell number estimates, see above).

For proliferation analysis, confocal stacks were acquired from tissue stained for Sox9 and Ki67. Sox9+ astrocytes were counted first and then assessed for Ki67 labeling. Cells that were Sox9+Ki67+ were counted and then analyzed for Cre activity based on GFAP-tdT signal to determine if Cre+ and Cre-cells express Ki67 at different rates in *Tmem30a^aKO^* animals.

For apoptosis analysis, confocal stacks were acquired from tissue stained for Sox9 and cleaved caspase-3 (CC3). Individual Sox9+ cells were not counted due to the extremely low rates of CC3 double labeling (Puñal et al., 2019). Instead, we assessed the number of CC3+Sox9+ double labeled cells within 12 confocal Z-stacks per retina (n = 2-3 per genotype), acquired as described above, with XY dimensions of 212.13 µm x 212.13 µm. These images sampled from all retinal eccentricities. At P5, each field of view contained approximately 125 astrocytes in control animals and 100 astrocytes in mutant animals (based on average Sox9 counts of similar images acquired for the analysis shown in Figure 2F). Therefore, approximately 6,500 astrocytes were analyzed across 60 fields of view and only 1 astrocyte was found to be CC3+ (Supplemental Figure 2F,G).

#### Quantification of cholinergic neurons

For *Tmem30a^sacKO^* analysis, at least 5 60x images were collected from each retina. Images were collected only from sections that were truly cross-sectional, perpendicular to the retinal layers, and containing the optic nerve head. ON and OFF starburst amacrine cells (SACs) were counted separately but combined for analysis because there was no difference in their density across *Tmem30a* genotypes.

#### Quantification of electroporated cortical astrocyte and radial glia density

For the cell counting assays in PALE brains, tilescan images of the whole injected hemisphere were acquired for each analyzed brain section on a Nikon A1 confocal microscope using the resonant scanner and 20x objective. The analysis was done on 2-3 sections evenly spaced through the electroporated region. Stitched images were created using NIS-Elements software. For each brain section, all PALE-tdT+ cortical astrocytes were counted. A few tdT+ neurons (presumably late-born) were characterized by their neuron-like morphology and excluded from this analysis. After counting, the analyzed area was measured using the Polygon selection tool in Fiji to determine astrocyte density (cells/mm^2^). For statistical analysis, densities were averaged across the 2-3 hemispheres that were analyzed for each animal.

To show that the difference in astrocyte density between genotypes was not due to differences in electroporation efficiency between the two groups, radial glia numbers were analyzed as an electroporation control. A 1mm-wide ROI, chosen to capture the highest density of PALE-tdT+ radial glia, was selected for analysis in each P7 hemisphere. Using a maximum-intensity Z-projection of the analysis region, each radial glia process that traversed at least half of the cortical depth was manually counted. Total radial glia count was divided by the width of the analyzed area to find their density per millimeter of subventricular zone length.

#### Astrocyte and vessel network analysis

For quantification of astrocyte and vessel network coverage, confocal Z-stacks spanning the entire depth of the RNFL and GCL were collected from GFAP- and/or Lectin-stained retinal wholemounts. Images sampled all eccentricities of the retina as described above. Using Fiji software, stacks were maximum-intensity-projected and GFAP+ or Lectin+ pixels were segmented using automatic thresholding (settings: Li, black and white, dark background). Total network coverage was then quantified by measuring the percentage of GFAP+ or Lectin+ pixel area in each image. To make this calculation, the “Analyze Particles” function in Fiji was used to quantify the area of segmented/binarized signal (settings: size 0–Infinity, analyzed in pixel units, circularity 0.00–1.00). Average percent area coverage was calculated using measurements from across the retina and is not weighted by eccentricity.

#### Software

All flow cytometry analysis and compensation matrices were performed in FlowJo software. All images were collected on a Nikon A1 confocal using NIS-Elements software or an Olympus Fluoview FV3000 using FV31S-SW software. All image processing was completed using Fiji software (Schindelin et al., 2012). All statistical analyses were performed and all graphs were made in GraphPad Prism software.

## ACKNOWLEDGEMENTS

This work was supported by the National Eye Institute (EY030611 to J.N.K; EY035853 to S. Gerecht, J.N.K. co-investigator; EY035254 to M. Samuel, J.N.K. co-investigator; EY034134 to X.C.; EY5722 to Duke University); Research to Prevent Blindness (Stein Innovation Award to J.N.K.; unrestricted grant to Duke University); The Kim B. and Stephen E. Bepler Endowment (S.C.F.); and a Duke Holland-Trice Award (graduate student fellowship award to C.E.P.). C.E. is an HHMI Investigator. We thank Wellcome Trust Sanger Institute for providing the mutant mouse line (Allele: *Tmem30a^tm1a(KOMP)Wtsi^*); INFRAFRONTIER/EMMA (www.infrafrontier.eu), and the EMMA node at the Institute of Molecular Genetics from which the mouse line was distributed (RRID:IMSR_EM:09580). We thank Joan Kalnitsky and Oleg Kolupaev in the Duke Eye Center Flow Cytometry core for providing access to and expertise with cell sorters; Ariane Pendragon for mouse colony management; and Alexandra Barker and Sarah Hadyniak for their comments on the manuscript.

## Key Resources Table

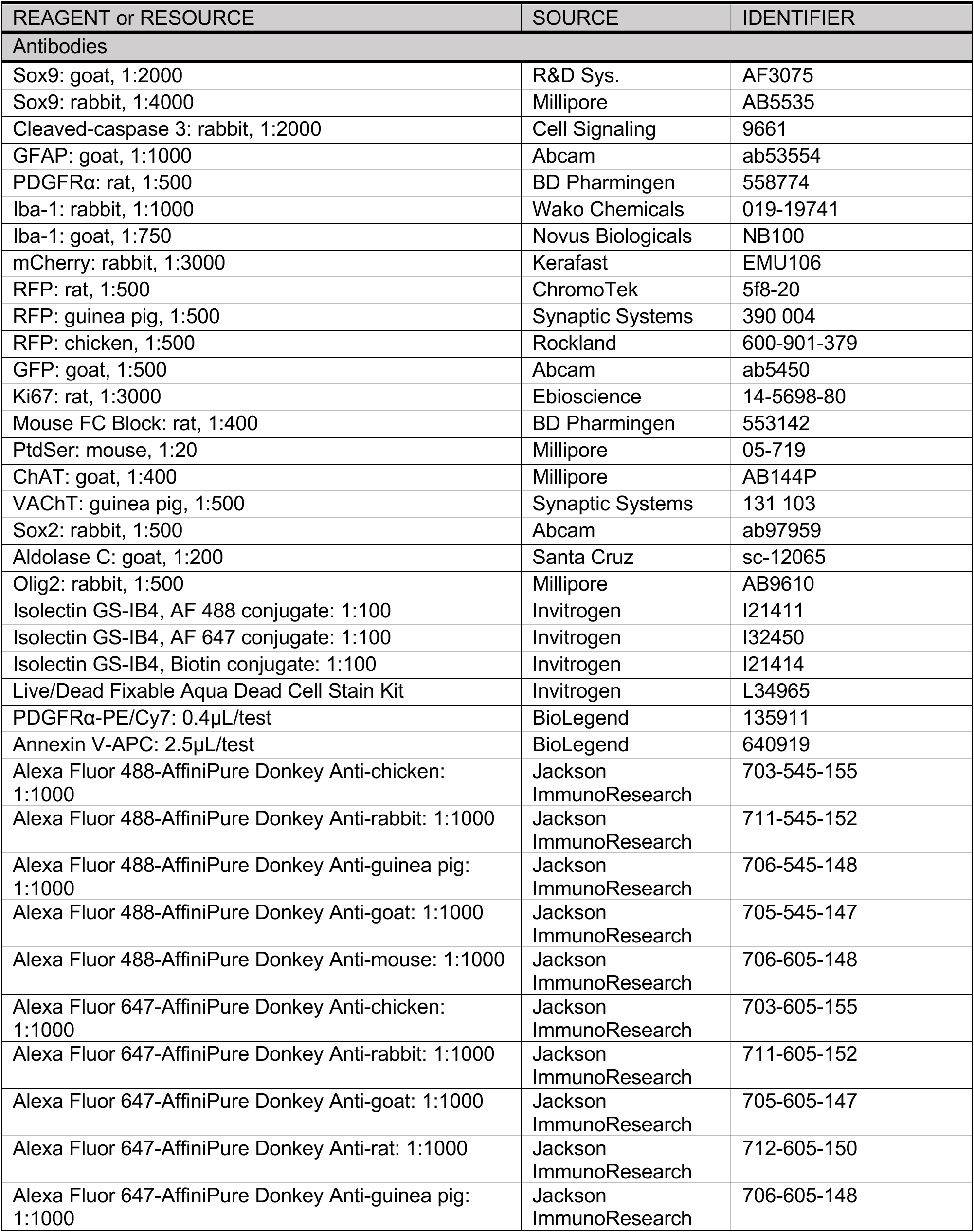
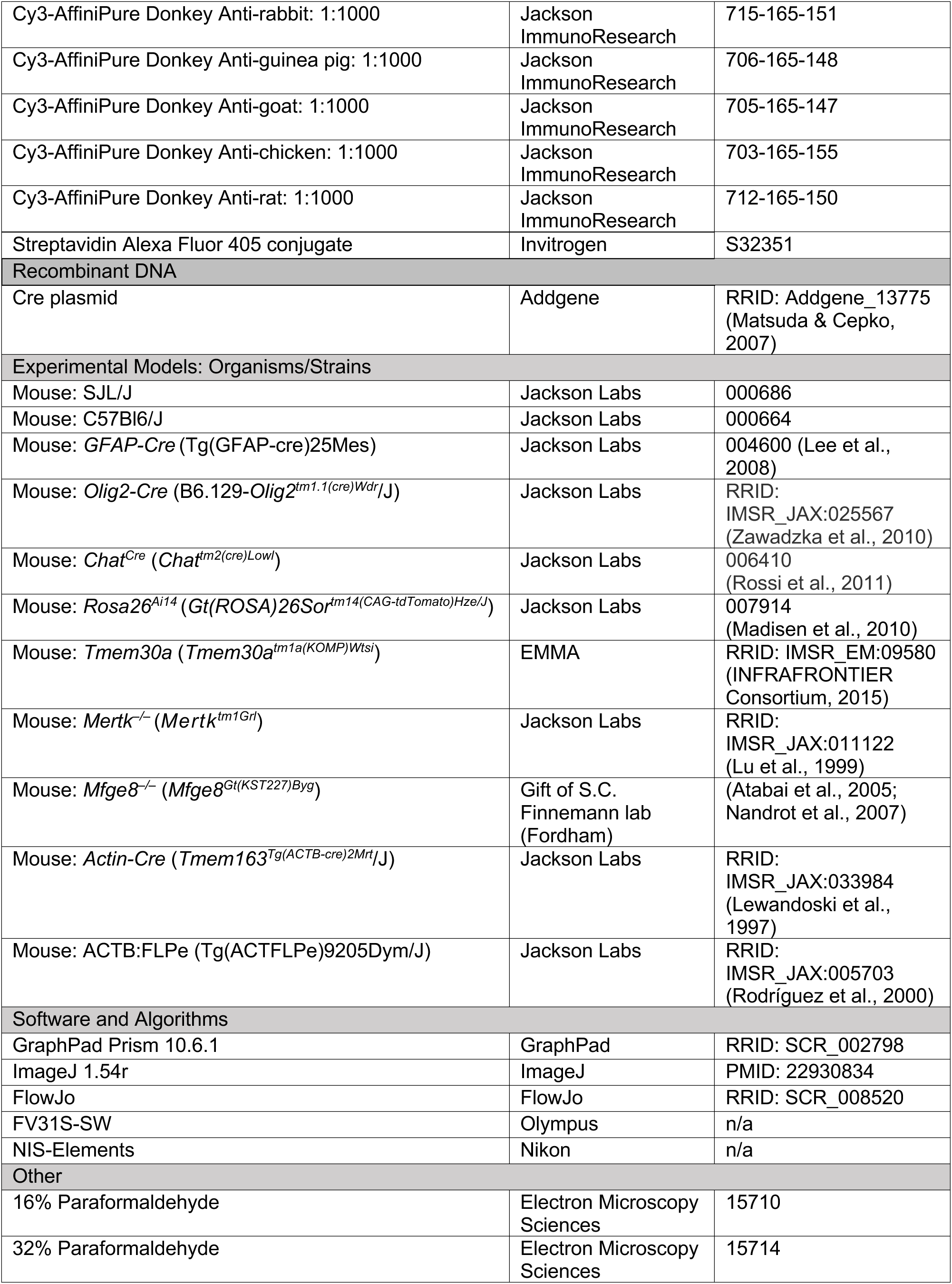
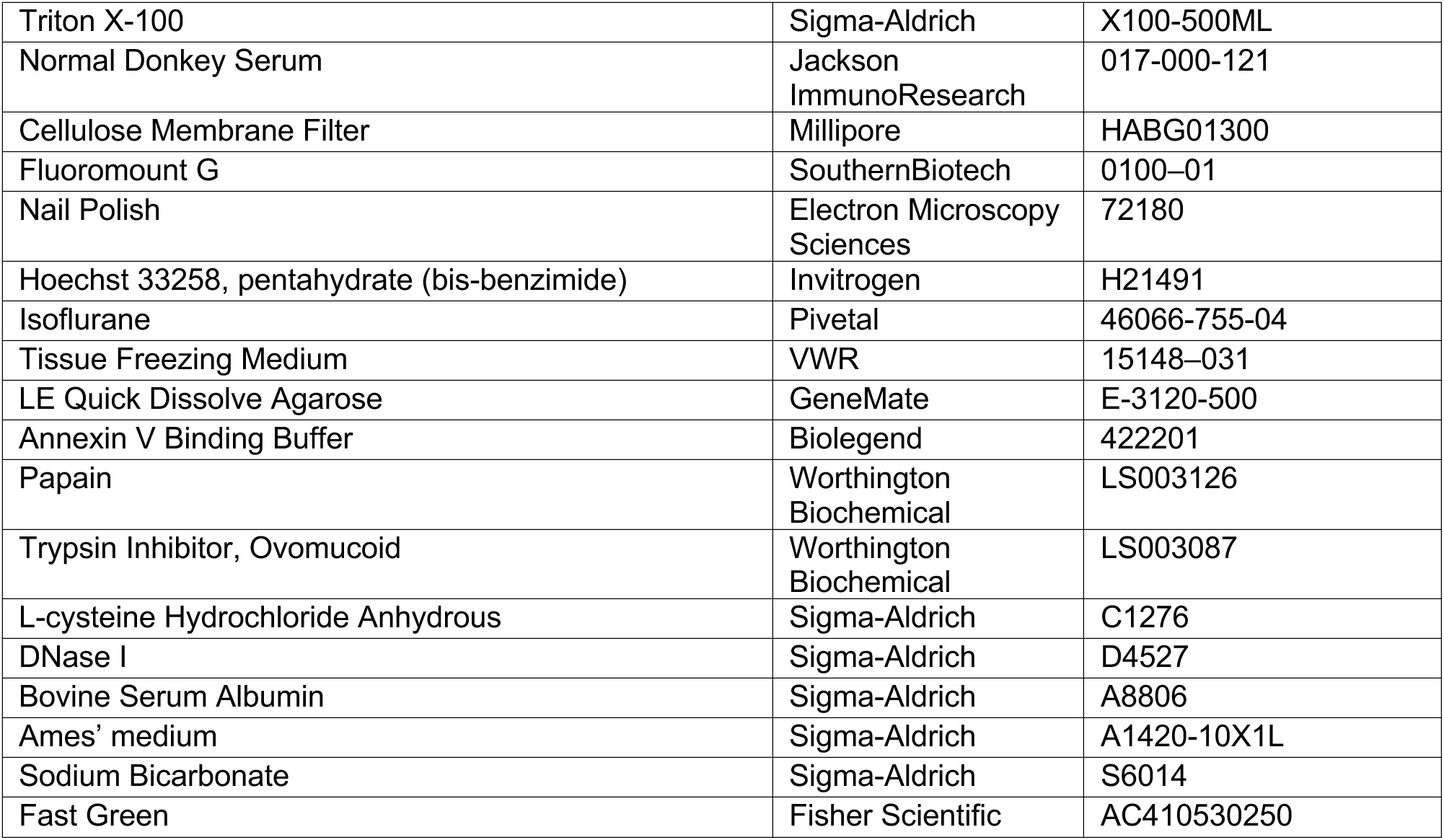

## Supplemental Figures

**Supplemental Figure 1:**
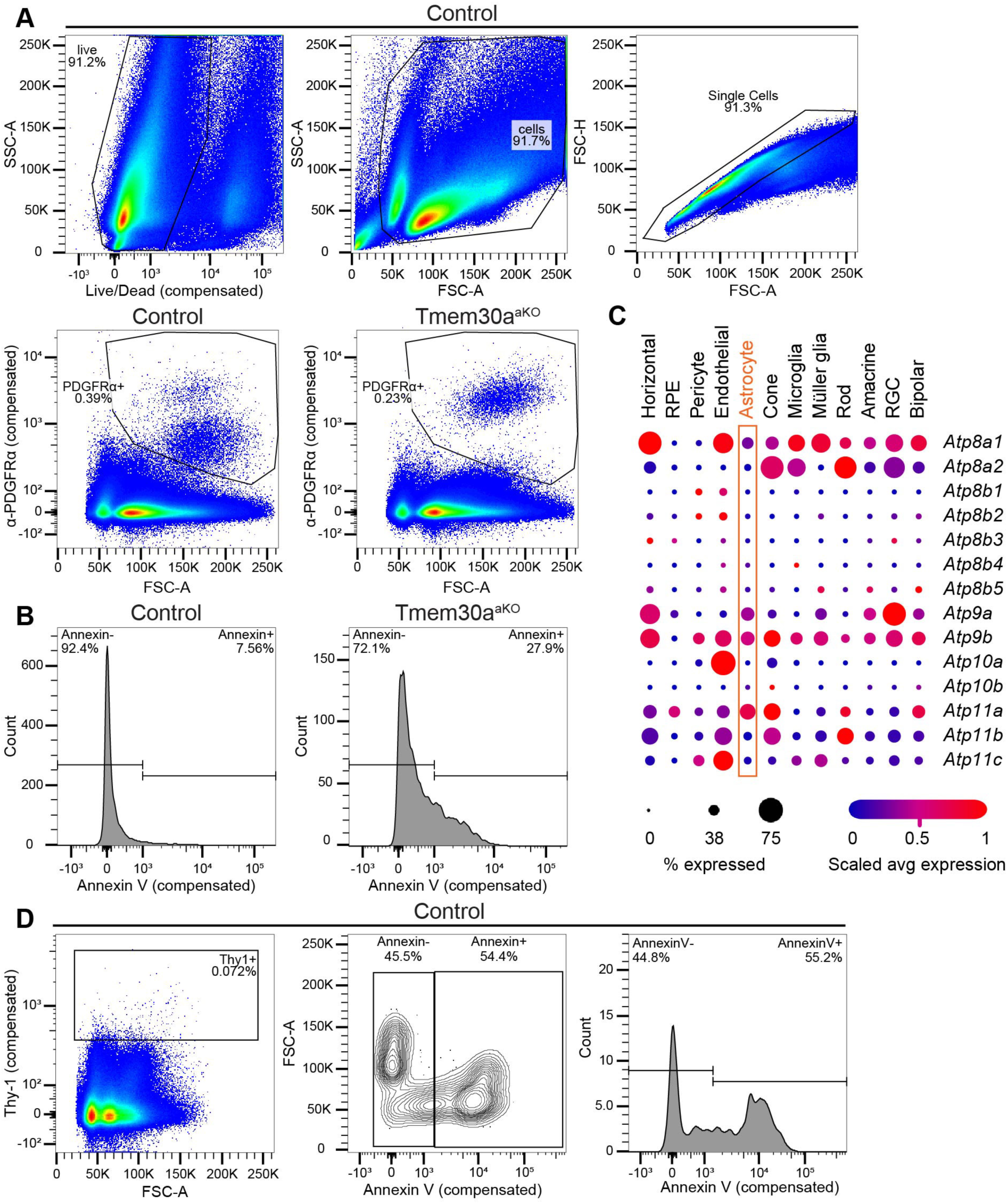
Gating strategy for Annexin V flow cytometry analysis. A) Top row: Three preliminary gates were drawn to exclude dead cells (left), cell debris (center), and doublets (right). Bottom row: Gates used for isolation of the PDGFRα+ astrocyte population. Control mice (left) and *Tmem30a^aKO^* mice (right) from the same litter are shown here. Cells that passed all of these gates were used to generate the plots in Figure 1C,D. Note the presence of a low and a high PDGFRα population in control animals, whereas *Tmem30a^aKO^* astrocytes mainly belonged to the high-PDGFRα population. This likely reflects a delay in astrocyte maturation, as PDGFRα is more highly expressed by astrocyte precursor cells than mature astrocytes (see Figure 1A). B) Gating strategy to determine the Annexin V+ and Annexin V– populations of PDGFRα+ astrocytes (i.e. cells passing gates shown in A). C) Dot plot showing expression of the 14 genes comprising the P4-ATPase gene family, as assessed in the Mouse Retinal Cell Atlas single-cell RNA-sequencing dataset (Li et al., 2024). Multiple members of this family are expressed by astrocytes (orange box). D) Strategy to validate the choice of Annexin V fluorescence cutoff used to draw the gates shown in B. Because there was no clear separation between Annexin V+ and Annexin V–populations within astrocytes or total retinal cells, we used a different cell type – Thy1+ cells (i.e. retinal ganglion cells) to determine where the cutoff between Annexin V+ and Annexin V–populations should be placed. Left; gate for Thy1+ cells (starting from cells that passed the gates shown in the top row of A. Center, right: two different plots showing a clear separation between Annexin V+ (likely apoptotic) and Annexin V– Thy1 populations. The Annexin V fluorescence cutoff shown in center, right plots was used as the cutoff for all astrocyte studies.

**Supplemental Figure 2:**
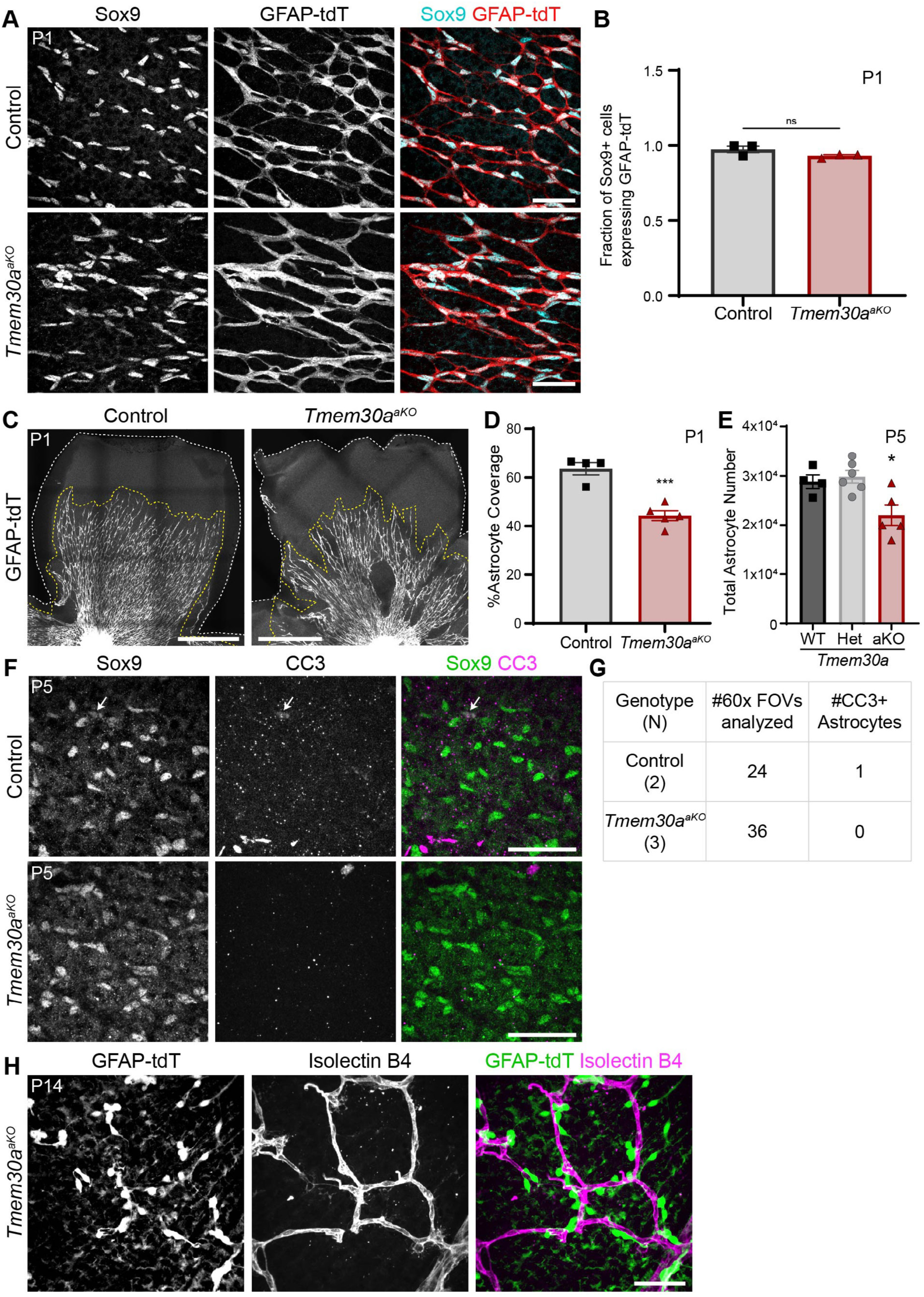
Additional phenotyping of *Tmem30a^aKO^* mice. A,B) Validation of retinal astrocyte Cre expression pattern in the GFAP-Cre mouse line. A: Representative images of P1 control and *Tmem30a^aKO^*retinas showing near-complete overlap of Sox9 (cyan) and GFAP-tdT (red) expression. B: Quantification of overlap between Sox9 and GFAP-tdT in P1 images similar to A. In control retina, 97% of Sox9+ astrocytes were tdT+, showing that recombination is nearly universal across the astrocyte population already by P1. In *Tmem30a^aKO^* retina, 93% of astrocytes were tdT+, showing that the preferential killing of Cre+ astrocytes is not seen this early in development. Statistics: two-tailed *t* test (not significant, *p* = 0.139). C,D) At P1, a smaller area of the retina is covered by astrocytes in *Tmem30a^aKO^*retinas, suggesting a delay in astrocyte migration. C: Representative images of P1 control and *Tmem30a^aKO^* retinas, showing astrocytes labeled with GFAP-tdT. White dotted lines delineate edge of retinal tissue. Yellow dotted lines delineate astrocyte wavefront. D: Quantification of astrocyte coverage of the retina, from images similar to C. For each retina, total area of astrocyte coverage (C, yellow line) was divided by total retina area (C, white line) to determine this percentage. Astrocyte coverage is significantly lower in *Tmem30a^aKO^* retinas compared to controls (44% vs. 64%). Statistics: two-tailed *t* test, \*\*\**p* = 4.8 x 10^-4^. E) Total astrocyte numbers at P5. Control animals plotted in Figure 2F have been separated into *Tmem30a^+/+^* and *Tmem30a^fl/+^* groups to show there is no phenotypic difference between the two genotypes. Statistics: one-way ANOVA showing a significant group effect (\**p* = 0.009), with post-hoc Tukey’s tests corrected for multiple comparisons. No significant difference between Tmem30a wildtype and heterozygotic animals (*p* = 0.905). F,G) Retinal astrocytes (Sox9+) generally do not express cleaved caspase 3 (CC3), a marker of apoptotic cells. F: Example images from P5 control and *Tmem30a^aKO^* retinas showing colocalization between Sox9 and CC3. G: Table showing the number of 60x fields of view analyzed in both control and *Tmem30a^aKO^* retinas and the number of CC3+ astrocytes found in this analysis. CC3+ cells were almost entirely Sox9-negative, indicating that they are not astrocytes. Control image in F was chosen to show the sole CC3+ astrocyte found in this analysis. H) Example image showing that the majority of surviving Cre+ astrocytes (GFAP-tdT, green) in a P14 *Tmem30a^aKO^* retina are closely associated with vessels (Isolectin B4, magenta) in central retina. This close association suggests that astrocytic interaction with vessels may promote cell survival even when PtdSer exposure is dysregulated in these cells. Error bars: mean ± SEM. Scale bars: A,E,G: 50 µm; C = 500 µm.

**Supplemental Figure 3:**
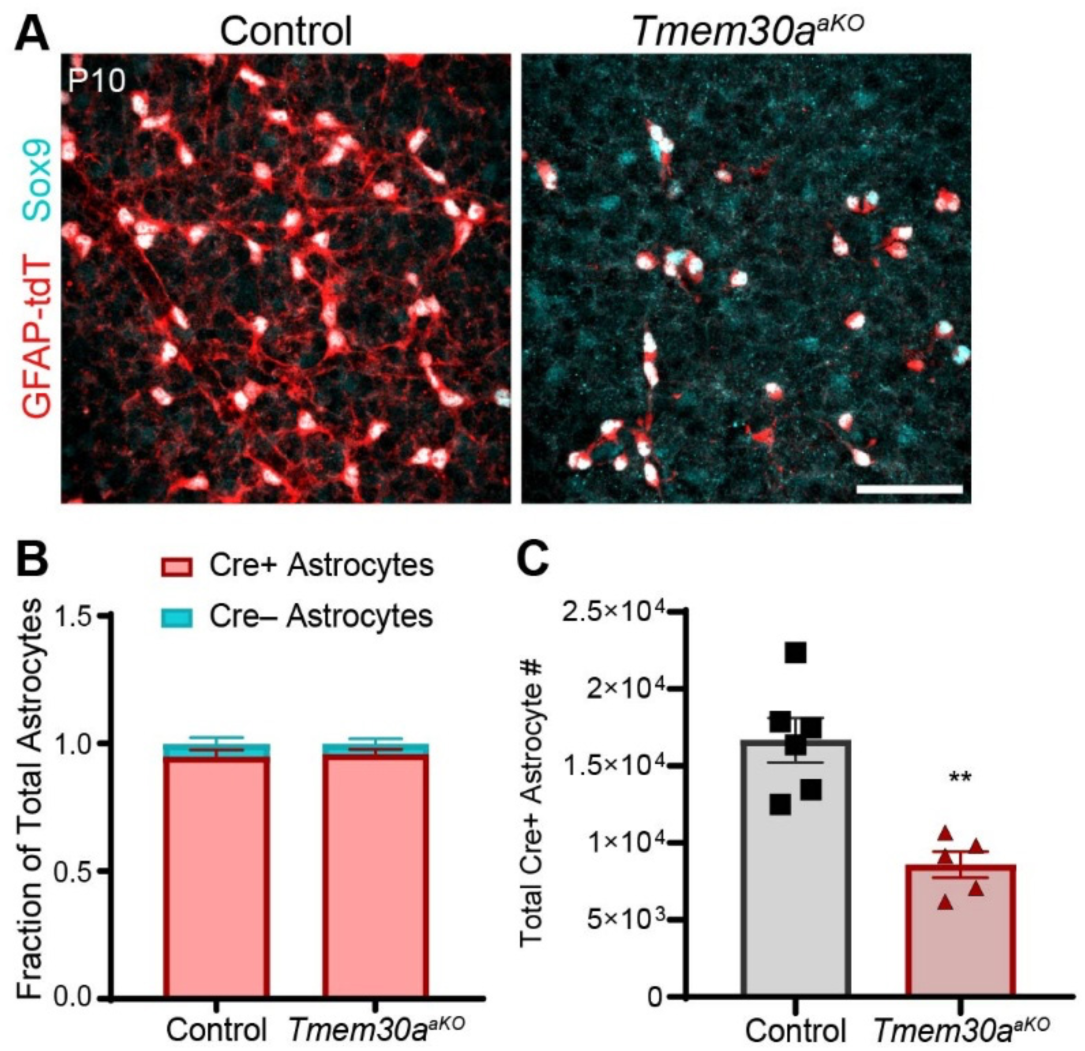
Astrocytes that were Cre-negative at P5 in *Tmem30a^aKO^* mice activate Cre expression by P10. A) Representative images of Sox9 (cyan) and GFAP-tdT (red) colocalization in P10 control and *Tmem30a^aKO^* retinas. B) Quantification of Cre+ (Sox9+GFAP-tdT+) and Cre– (Sox9+GFAP-tdT–) astrocytes at P10. Nearly all of the astrocytes are Cre+ for both control and *Tmem30a^aKO^* retinas (96% and 95%, respectively). Statistics: two-way ANOVA showing a non-significant interaction between genotype and colocalization (*p* = 0.7028523). C) Total Cre+ astrocyte numbers at P10, estimated from eccentricity-weighted density measurements and retina area. In contrast to the situation at P5 (Figure 3C-E), counting Cre+ astrocytes (this graph) and counting all Sox9+ astrocytes (Figure 2F) both yielded similar estimates of astrocyte cell number declines in *Tmem30a^aKO^* retinas. Statistics: two-tailed *t* test, \*\**p* = 0.001. Error bars: mean ± SEM. Scale bar: 50 µm.

**Supplemental Figure 4:**
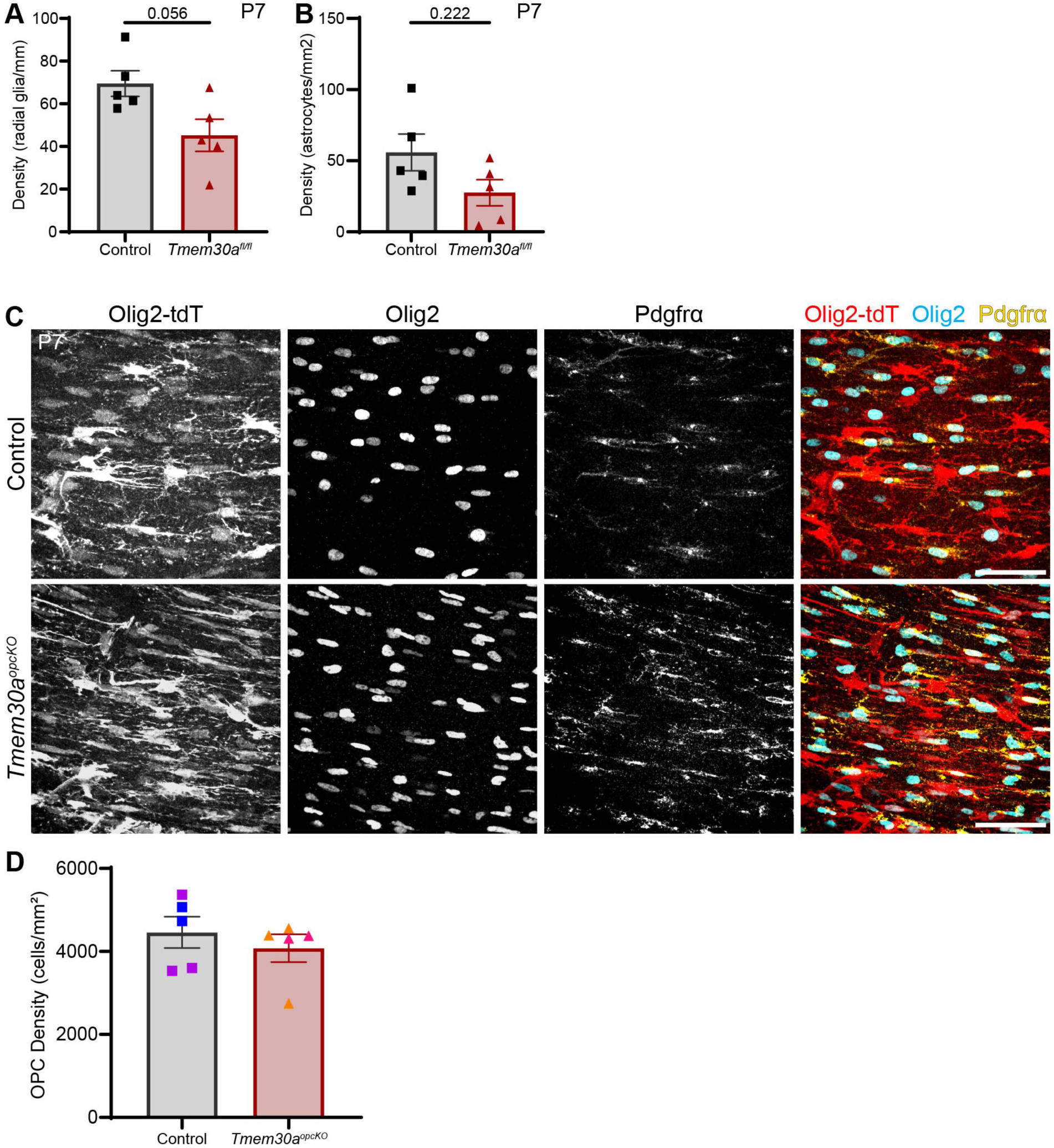
Loss of *Tmem30a* from brain glia. A,B) Replotting of radial glia and astrocyte densities from P7 PALE brain sections, which were shown in Figure 4D as a scatterplot but here are shown as individual graphs. Radial glia density was quantified as processes/mm of subventricular zone linear distance. Astrocyte density was quantified as astrocytes/mm^2^ of cortical area. Statistics: Mann-Whitney tests; *p* values shown on graphs. C,D) Analysis of OPC survival in *Tmem30a^opcKO^* mice. C: Representative images of corpus callosum white matter from P7 control and *Tmem30a^opcKO^*brain sections. D: Quantification of Olig2+ cell density in P7 control and *Tmem30a^opcKO^*brain sections. In images (C), *Olig2^Cre^*-driven tdTomato (Olig2-tdT, red) mostly overlaps with Olig2 (cyan) and PDGFRα (yellow) labeling, showing that the Cre+ cells in this mouse line are mostly oligodendrocyte precursor cells or mature oligodendrocytes. (Consistent with prior studies, we also observed a subset of tdT+ cells that were Olig2-negative, which are likely the subset of fibrous and protoplasmic astrocytes described by Zawadzka et al, 2010). No obvious change in *Tmem30a^opcKO^*cell density was observed; this impression was confirmed by OPC cell counts (D). For D, at least 2 sections from 2 brains/genotype were analyzed. Data points that are the same color on the graph are from the same animal. Statistics: two-tailed *t* test, ns *p* = 0.470. Error bars: mean ± SEM. Scale bars: A: 250 µm; C: 50 µm.

**Supplemental Figure 5:**
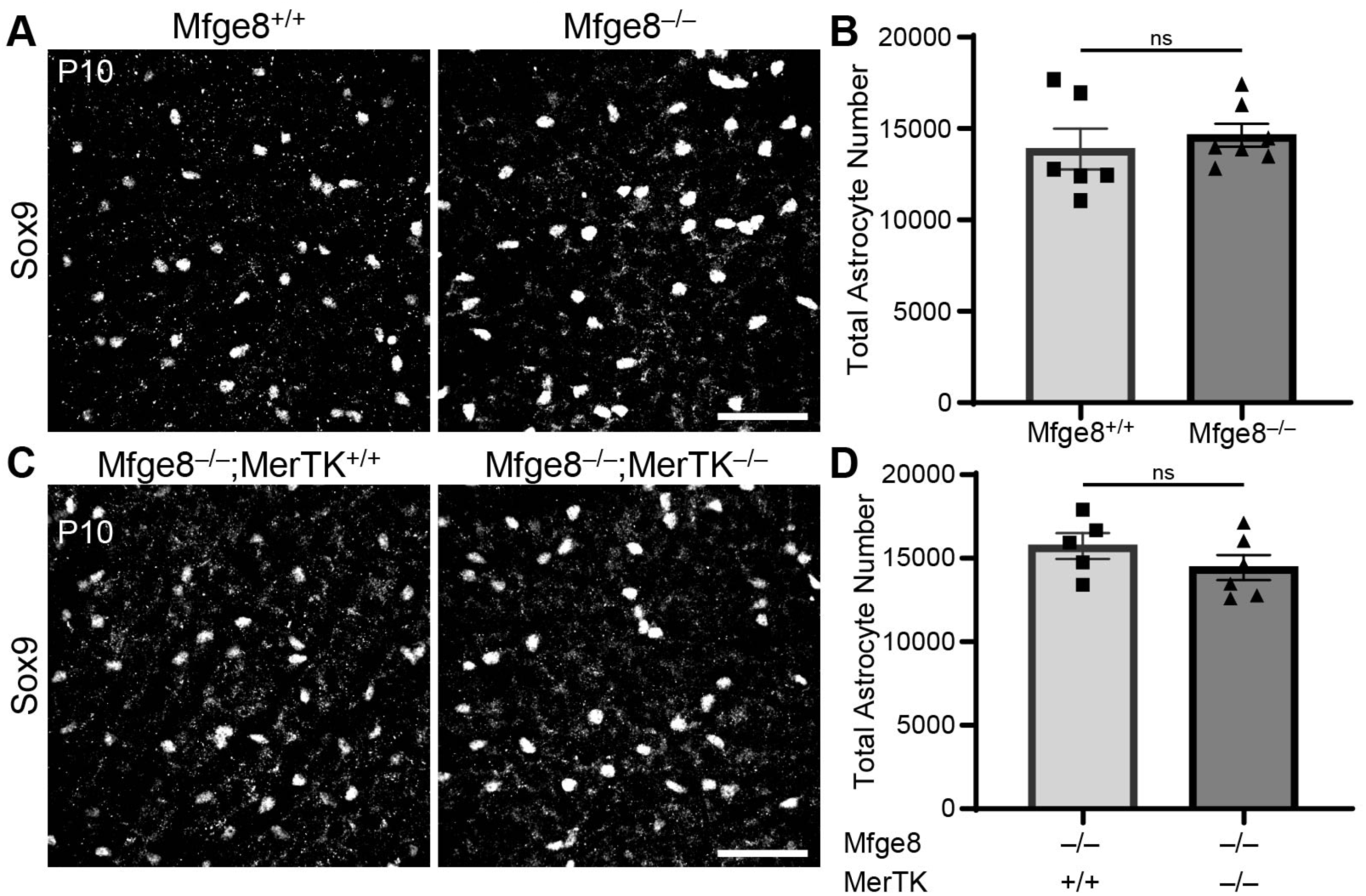
Neither *Mfge8* knockout nor *Mfge8; Mertk* double knockout affect astrocyte density or total number at P10. A,B) *Mfge8* single mutant analysis. A: Representative images of astrocytes labeled by Sox9 in central retina. B: Total astrocyte numbers in *Mfge8* wildtype and mutant mice at P10. Statistics: two-tailed *t* test, *p* = 0.554. C,D) *Mfge8;Mertk* double mutant analysis. C: Representative images of astrocytes labeled by Sox9 in central retina. D: Total astrocyte numbers at P10 in *Mfge8* single mutant animals and *Mfge8; Mertk* double mutant animals. Note that this is a different cohort of *Mfge8* single mutants than from the analysis shown in B. Statistics: two-tailed *t* test, *p* = 0.262. Error bars: mean ± SEM. Scale bars: 50 μm.

## Notes

### Competing Interest Statement

The authors have declared no competing interest.

